# Opponent vesicular transporters regulate the strength of glutamatergic neurotransmission in a *C. elegans* sensory circuit

**DOI:** 10.1101/2021.03.20.436253

**Authors:** Jung-Hwan Choi, Lauren Bayer Horowitz, Niels Ringstad

## Abstract

At chemical synapses, neurotransmitters are packaged into synaptic vesicles that release their contents in response to depolarization. Despite its central role in synaptic function, regulation of the machinery that loads vesicles with neurotransmitters remains poorly understood. We find that synaptic glutamate signaling in a *C. elegans* chemosensory circuit is regulated by antagonistic interactions between the canonical vesicular glutamate transporter EAT-4/VGLUT and another vesicular transporter, VST-1. Loss of VST-1 strongly potentiates glutamate release from chemosensory BAG neurons and disrupts chemotaxis behavior. Analysis of the circuitry downstream of BAG neurons shows that excess glutamate release disrupts behavior by inappropriately recruiting RIA interneurons to the BAG-associated chemotaxis circuit. Our data indicate that *in vivo* the strength of glutamatergic synapses is controlled by regulation of neurotransmitter packaging into synaptic vesicles via functional coupling of VGLUT and VST-1.

## INTRODUCTION

Flow of information through neural circuits requires the regulated release of neurotransmitters at chemical synapses. The neurotransmitters that mediate synaptic signaling are stored in synaptic vesicles. These remarkable organelles accumulate neurotransmitters above their cytoplasmic concentrations and fuse with the plasma membrane upon neuronal depolarization to release their contents into the synaptic cleft. This fusion mechanism is common to neurons of all types^1, 2^. By contrast, the mechanisms that package neurotransmitters into synaptic vesicles vary according to the neurotransmitter identity of a neuron. Each neurotransmitter is associated with a vesicular transporter that mediates its influx into the vesicular lumen^3^. Distinct transporters support the loading of synaptic vesicles with glutamate, GABA, acetylcholine, and monoamines such as serotonin and dopamine. Consequently, vesicular transporters are determinants of a key functional attribute of every neuron: what neurotransmitters that neuron uses to signal to synaptic partners.

Neurons can express more than one vesicular transporter, and this not only determines which transmitters are released but also might determine how much of each transmitter is packaged into synaptic vesicles. Because all vesicular transmitters harness the energy stored in an electrochemical gradient generated by a vesicular proton pump (reviewed in ^4^), one vesicular transporter can influence the function of another. Vesicular glutamate transporter (VGLUT) 3 and vesicular acetylcholine transporter (VAChT) are co-expressed in striatal cholinergic interneurons, and glutamate transport through VGLUT3 potentiates acetylcholine transport by VAChT by allowing the vesicular proton pump to generate a steeper pH gradient across the vesicle membrane^5^. Similarly, glutamate transport via VGLUT2, which is co-expressed in a subset of neurons with the vesicular monoamine transporter (VMAT), enhances VMAT-dependent dopamine transport into vesicles^6, 7^. Based largely on biophysical studies, it has been suggested that interactions between vesicular transporters can regulate the strength of neurochemical signaling in specific circuits. The functional significance of interactions between vesicular transporters in neural circuits and on behavior, however, remains poorly understood. It is also likely that these interactions are widespread and involve additional vesicular transporters and channels, such as cation/H^+^ exchangers and ClC chloride channels^8–, 11^, which do not transport neurotransmitters but mediate the transport of other ions that can affect the electrochemical gradient. New vesicular transporters continue to be discovered^12, 13^, indicating that the current census of vesicular transporters is incomplete and raising the intriguing possibility that many neurotransmitter systems are regulated by interactions between vesicular transporters.

The nematode *C. elegans* is a powerful model for studying synaptic function in general and vesicular transporters in particular. Genetic and biochemical studies of *C. elegans* revealed the molecular identity of VAChT^14^ and VGAT (vesicular GABA transporter)^15^. The characterization of the primary *C. elegans* VGLUT, EAT-4, and the assignment of its function to glutamatergic neurons in the pharyngeal nervous system helped establish the molecular identities of VGLUTs^16, 17^. A particular strength of the *C. elegans* model is that simple and stereotyped behaviors critically depend on specific neurotransmitter signals. Genetic analysis of such behavior is a powerful method to identify molecular factors required for specific kinds of neurotransmission.

Synaptic glutamate signaling in *C. elegans* is required for a suite of chemosensory, mechanosensory, and thermosensory behaviors^18, 19^. As in the vertebrate brain, glutamate is a major excitatory neurotransmitter in the *C. elegans* nervous system^16, 20, 21^. Of note, the molecular mechanisms of glutamatergic neurotransmission are highly conserved between *C. elegans* and vertebrates. The *C. elegans* nervous system uses homologs of ionotropic AMPA receptors, kainate receptors, and NMDA receptors for fast, excitatory synaptic signaling^22–25^. Homologs of metabotropic glutamate receptors mediate G protein-coupled glutamate signaling in the *C. elegans* nervous system^26–29^. *C. elegans* neurons and glia use conserved mechanisms to package glutamate into synaptic vesicles and clear glutamate after its release, respectively^16, 30, 31^.

Here, we report the discovery of a vesicular transporter, VST-1, that is required in glutamatergic chemosensory neurons for chemotactic avoidance behavior. Loss of VST-1 causes a dramatic increase in the amount of glutamate released from CO_2_-sensing neurons, which requires EAT-4/VGLUT. We find that excess glutamate signaling in *vst-1* mutants recruits interneurons to the chemotaxis circuit that are normally quiescent via AMPA-type glutamate receptors, and that this ectopic activation of interneurons in *vst-1* mutants is a major cause of their chemotaxis defects. These data show that a presynaptic mechanism that antagonizes VGLUT-dependent packaging of glutamate into synaptic vesicles is a critical determinant of the strength of synaptic glutamate signaling *in vivo*.

## RESULTS

*C. elegans* possesses a pair of chemosensory neurons that detect the respiratory gas carbon dioxide (CO_2_), the BAG neurons. When we analyzed their transcriptome for differentially expressed genes^32^ homologous to known vesicular transporters, we observed that BAG neurons are enriched for transcripts encoding EAT-4, the primary *C. elegans* VGLUT (**Figure 1a, c**). We also noted that they are enriched for transcripts that encode the related transporter SLC-17.1 (**Figure 1b, c**). As indicated by its name, SLC-17.1 is a member of the SLC17 family of solute transporters, which comprises the VGLUTs, the vesicular nucleotide transporter (VNUT), sialin, and inorganic phosphate transporters^33^ (**Figure 1d**). For clarity, we hereafter refer to *slc-17.1* as *vst-1* (*vst* = *<u>v</u>esicular <u>s</u>olute transporter*).

**Figure 1.**
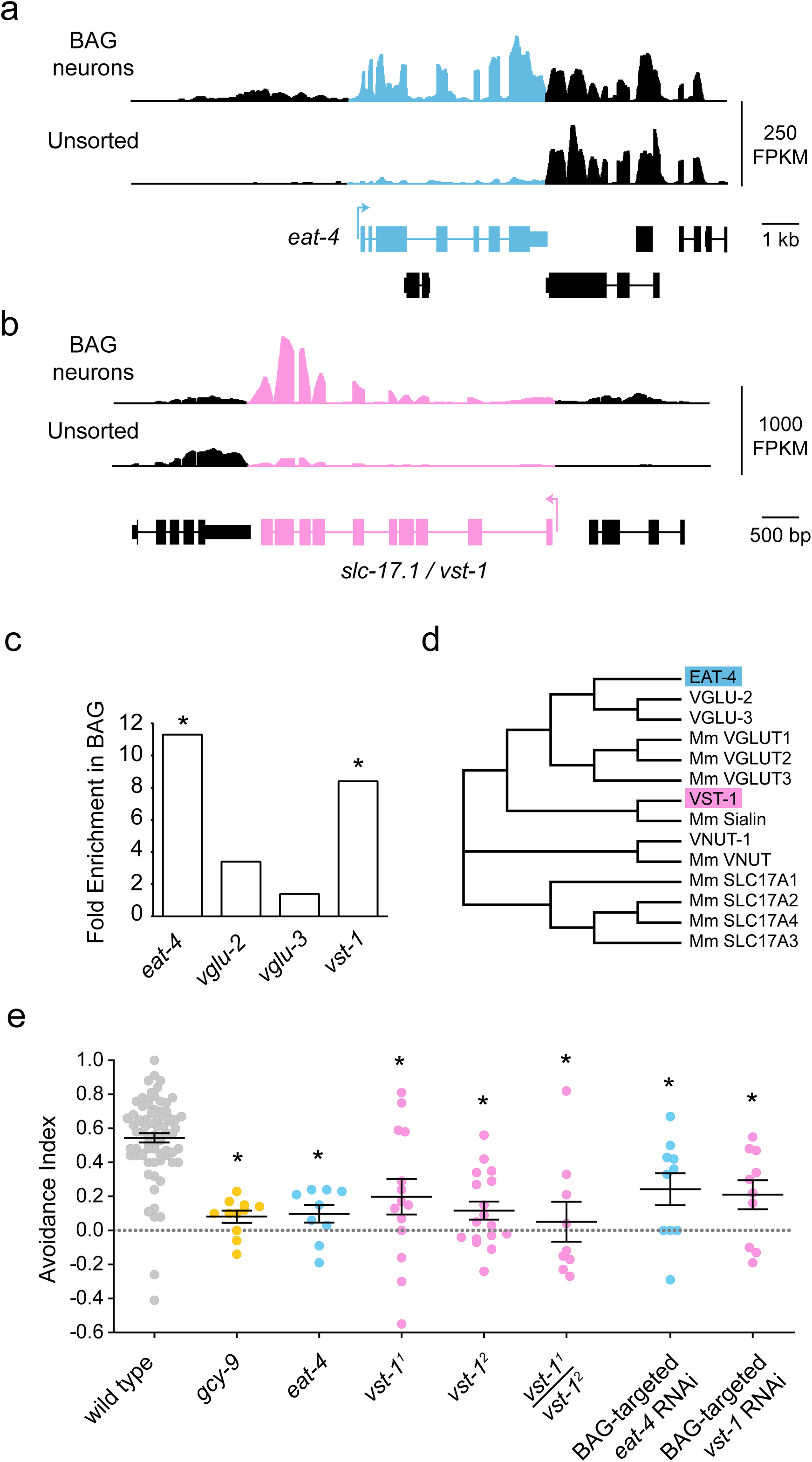
*vst-1* is required in BAG chemosensory neurons for CO_2_ chemotaxis. (**a**) Mean RNAseq signal (Fragments Per Kilobase of transcript per Million mapped read, FPKM) of *eat-4* obtained from two biological replicates. *eat-4* exons are blue and exons from neighboring genes are black. (**b**) Mean RNAseq signal (FPKM) of *vst-1* (*slc-17.1*) obtained from two biological replicates. *vst-1* exons are red and exons from neighboring genes are black. (**c**) Fold enrichment in BAG neurons of *eat-4* transcripts and transcripts from *eat-4*-like genes. Asterisks indicate false discovery rates of less than 0.01. (**d**) Dendrogram of a subset of *C. elegans* SLC17 family transporters related to VST-1 (species not indicated) and *M. musculus* SLC17 family transporters (indicated by Mm). SLC17A1-4 indicate the type I sodium-dependent inorganic phosphate transporters of the family. (**e**) Avoidance indices (mean ± SEM) of *vst-1* and *eat-4* mutants from measurements of CO_2_-chemotaxis (*n* = 74, 10, 9, 14, 17, 9, 10, and 10 for wild type, *gcy-9*, *eat-4*, *vst-1^1^*, *vst-1^2^*, *vst-1^1^*/*vst-1^2^*, BAG-targeted *eat-4* RNAi, and BAG-targeted *vst- 1* RNAi, respectively). Asterisks indicate p < 0.05 calculated from comparisons of the mutants to wild type using a Kruskal-Wallis test corrected for multiple comparisons with Dunn’s test (p-values for each comparison marked with an asterisk - wild type vs *gcy-9(tm2816)*: <0.0001; wild type vs *eat-4(ky5)*: 0.0003; wild type vs *vst-1^1^*: 0.0040; wild type vs *vst-1^2^*: <0.0001; wild type vs *vst-1^1^/vst-1^2^*: 0.0002; wild type vs BAG-targeted *eat-4* RNAi: 0.026; wild type vs BAG-targeted *vst-1* RNAi: 0.0094). For simplicity, *vst-1^1^* was used to denote *vst-1(gk673717)* and *vst-1^2^* was used to denote *vst-1(gk308047)*.

To determine whether VST-1 is required for the function of chemosensory BAG neurons, we tested whether mutation of *vst-1* or *vst-1* knockdown by RNAi affected a chemotaxis behavior supported by BAGs. Under basal conditions, *C. elegans* avoid CO_2_ and navigate down a CO_2_ gradient^34, 35^. As expected, wild-type animals placed in an arena with sectors containing either CO_2_-enriched air or air with no CO_2_ avoided the CO_2_-enriched sector (**Figure 1e**). Animals lacking the CO_2_ receptor GCY-9^36, 37^ were profoundly defective in CO_2_-avoidance and partitioned equally between the two sectors (**Figure 1e**), consistent with previous studies^36, 38^. Animals lacking EAT-4/VGLUT were also severely defective in CO_2_-avoidance, indicating that glutamatergic signaling is essential for this behavior (**Figure 1e**). Mutants carrying nonsense alleles of *vst-1* (**Supplemental Figure 1a**) were also defective for CO_2_-avoidance, as were trans- heterozygotes for different nonsense alleles (**Figure 1e**). These data show that VST-1 is required for BAG-dependent chemotaxis.

To test whether *vst-1* functions in BAG neurons, we expressed double-stranded RNA corresponding to *vst-1* coding sequences specifically in BAGs to trigger RNAi and knock down *vst-1*. BAG-specific RNAi targeting *vst-1* caused a CO_2_-avoidance defect (**Figure 1e**), suggesting *vst-1* is required in BAG neurons. We further observed that the effect of *vst-1* knockdown in BAGs was comparable to that of knocking down *eat-4/VGLUT* (**Figure 1e**), indicating the functional importance of VST-1 in these glutamatergic sensory neurons.

### VST-1 is a synaptic vesicle transporter expressed in a majority of glutamatergic neurons

To determine the cellular expression pattern of *vst-1*, we generated a fosmid reporter that expresses nuclear mCherry in cells that express *vst-1* (**Supplemental Figure 1b**). We made transgenic animals carrying this reporter together with an *eat-4* fosmid reporter that expresses nuclear YFP in glutamatergic neurons^21^. These reporters indicated that *vst-1* is expressed in many head neurons (**Figure 2a**). Notably, a large majority of glutamatergic neurons in the head marked by the *eat-4* reporter also expressed the *vst-1* reporter (**Figure 2b**). We next asked where VST-1 localizes within neurons. For this, we generated another *vst-1* reporter that encodes a VST-1::GFP fusion (**Supplemental Figure 1c, d**). VST-1::GFP fluorescence was strikingly enriched in the nerve ring (**Figure 2c**), a neuropil containing most of the synaptic connections in the *C. elegans* nervous system. Many synaptic proteins, including components of synaptic vesicles, are expressed in a similar pattern^14, 15, 39^. To test whether VST-1 is associated with synaptic vesicles, we introduced this VST-1::GFP reporter into *unc-104* mutants, which lack a kinesin required for the transport of synaptic vesicle components, including vesicular transporters, to synapses^15, 40^. In *unc-104* mutants, VST::GFP was no longer highly enriched in the nerve ring and instead accumulated in neuronal cell bodies (**Figure 2c**). These data provide evidence that VST-1 is present on synaptic vesicles. To further confirm that VST-1 is associated with synaptic vesicles, we prepared VST-1::GFP transgenic animals for immunogold staining and electron microscopy. Ultrathin sections of the nerve ring showed the circular profiles of fasciculated axons, many of which contained clusters of synaptic vesicles that were marked with anti-GFP immunogold (**Figure 2d**). There was no immunogold labeling in the absence of anti-GFP primary antibody (**Supplemental Figure 2**), indicating that the immunogold label was specifically reporting VST-1::GFP localization. Together, these data indicate that VST- 1 is localized on synaptic vesicles and that it is expressed by a majority of glutamatergic neurons in the *C. elegans* nervous system.

**Figure 2.**
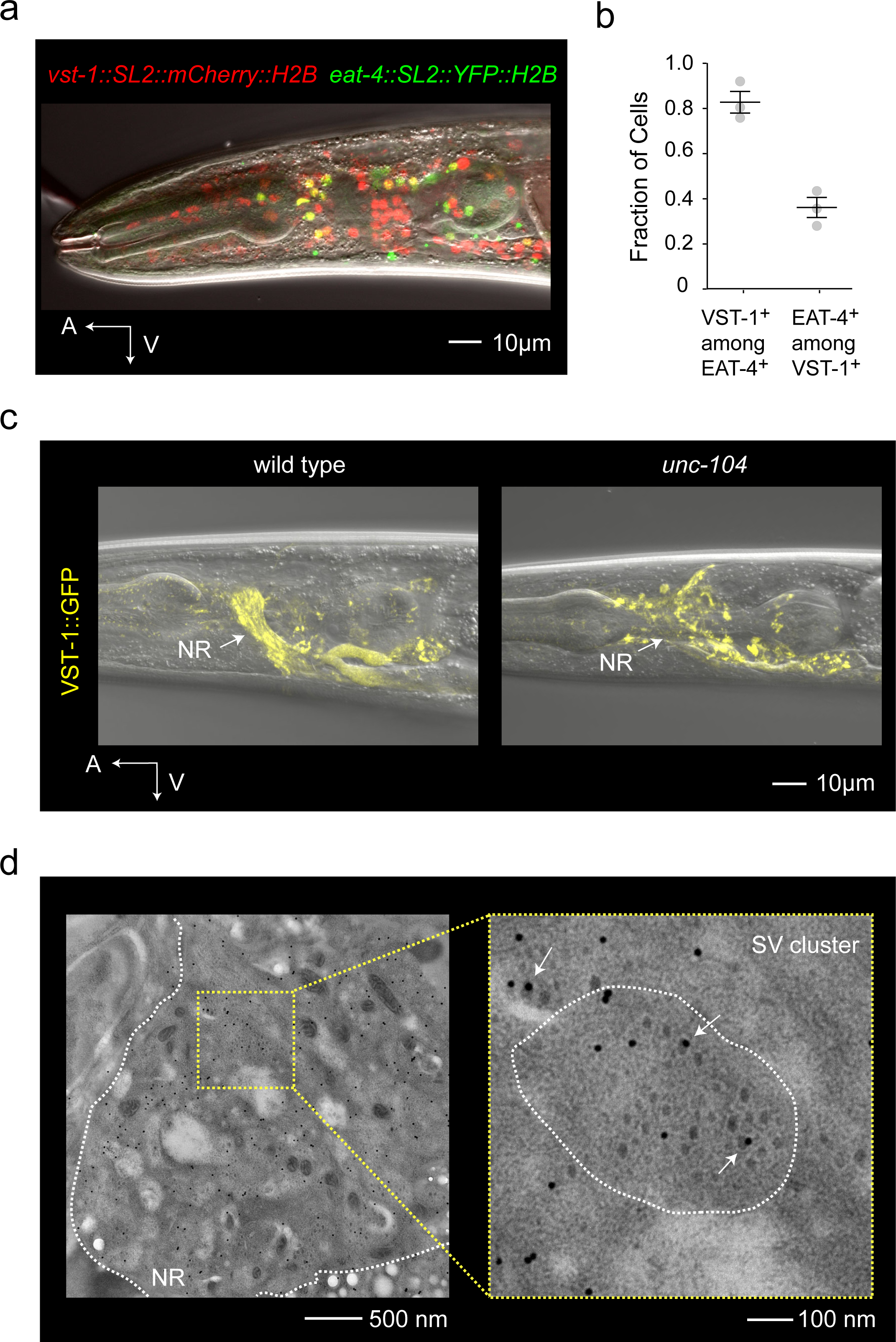
VST-1 is a synaptic vesicle transporter expressed by a majority of glutamatergic neurons. (**a**) Micrograph of an animal expressing fosmid reporters of *eat-4/vglut* and *vst-1*. The *eat-4* and *vst-1* reporters express nuclear YFP (pseudocolored green) and nuclear mCherry (red), respectively. (**b**) Quantification of *eat-4* and *vst-1* co-expression. The fraction of *eat-4*-expressing cells also expressing *vst-1* and vice-versa was determined from three independent experiments. (**c**) Localization of VST-1::GFP to presynaptic domains in the nervous system. The left panel shows a representative micrograph of a wild-type animal, in which VST-1::GFP is highly localized to the nerve ring (NR), marked with an arrow. The right panel shows VST-1::GFP expression in *unc-104/kinesin* mutants, which is required for proper transport of synaptic vesicle components into axons. (**d**) Immunoelectron micrographs of a strain expressing VST-1::GFP and stained with anti-GFP antibodies detected with 15 nm gold particles. The left panel shows a field of view at a magnification of 25000x. The boundary of the nerve ring is marked by a dashed white line. The right panel shows a magnified view of the region indicated by dashed yellow lines. The dashed white line in the right panel indicates the perimeter of a single neurite cut in cross-section. Immunogold particles (15 nm) show the expression of VST-1::GFP. Arrows indicate immunogold associated with synaptic vesicles.

### VST-1 inhibits EAT-4/VGLUT-dependent glutamate release from BAG neurons and can acidify synaptic vesicles

Because VST-1 is an SLC17-family transporter related to EAT-4/VGLUT, we hypothesized that VST-1 and EAT-4 might function together to mediate glutamatergic neurotransmission. To test this hypothesis, we designed an assay to measure evoked glutamate release from BAG neurons. We expressed the glutamate sensor iGluSnFR^30^ on the surface of BAG neurons, which we isolated in culture. Under these conditions, we could depolarize individual BAG neurons with high-potassium saline and measure changes in iGluSnFR fluorescence that reported glutamate release from that neuron. Wild-type BAG neurons displayed robust iGluSnFR signals upon depolarization (**Figure 3a**). iGluSNFR signals were severely diminished by CdCl_2_, a calcium channel inhibitor (**Supplemental Figure 3a, b**), indicating that these signals reflected calcium-dependent release of glutamate from BAG neurons. Also, BAG neurons lacking EAT-4/VGLUT did not display any evoked iGluSnFR signals (**Figure 3b**), further indicating that iGluSNFR signals report the evoked release of glutamate from BAG neurons. Of note, these data also failed to indicate any EAT-4/VGLUT- independent mechanism for glutamate release from BAGs. We next determined the effect of *vst-1* mutation on glutamate release from BAGs. Surprisingly, analysis of two independent *vst-1* mutants showed that loss of VST-1 significantly increased glutamate release from BAGs (**Figure 3c-f**). These data indicate that in BAG neurons VST-1 antagonizes EAT-4/VGLUT-dependent glutamate release.

**Figure 3.**
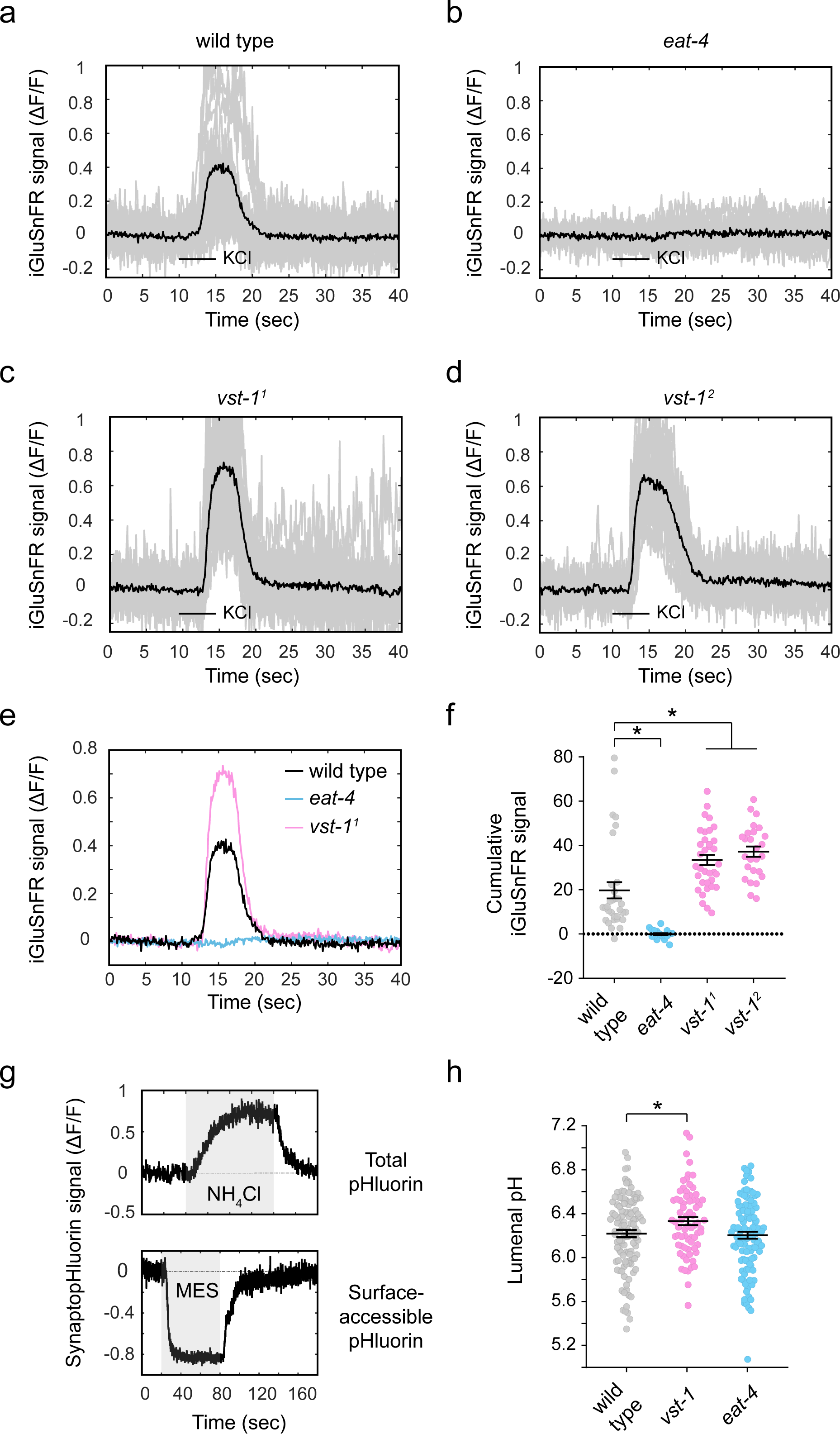
VST-1 inhibits EAT-4/VGLUT-dependent glutamate release from BAG neurons and acidifies synaptic vesicles. (**a-d**) iGluSnFR signal (ΔF/F) from neurites of BAG neurons in culture expressing the glutamate sensor iGluSnFR. Individual traces are plotted in gray and the mean signal is plotted in black. Neurons were depolarized with a five second pulse of 100 mM KCl. BAG neurons were isolated from the (**a**) wild type (*n* = 32), (**b**) *eat-4(ky5)* mutants (*n* = 19), (**c**) *vst-1(gk673717)* mutants (*vst-1^1^*) (*n* = 35), and (**d**) *vst-1(gk308047)* mutants (*vst-1^2^*) (*n* = 26). (**e**) Mean iGluSnFR signals from wild-type, *eat-4*, and *vst-1* BAG neurons. The overlay shows increased glutamate release from *vst-1* neurons and no glutamate release from *eat-4* neurons. (**f**) Quantification of iGluSnFR signals (mean ± SEM) from wild-type and mutant BAG neurons (*n* = 32, 19, 35, and 26 for wild type, *eat-4*, *vst-1^1^*, and *vst-1^2^*, respectively). Cumulative signal during between 10 and 20 sec was computed for each trial. Asterisks indicate *p* < 0.05 for comparisons of the mutants to wild type as determined by a Kruskal-Wallis test and corrected for multiple comparisons via Dunn’s test (p-values for each comparison marked with an asterisk – wild type vs *eat-4*: 0.0004; wild type vs *vst-1^1^*: 0.0035; wild type vs *vst-1^2^*: 0.0005). (**g**) Measurement of total synaptopHluorin and surface-accessible synaptopHluorin used to compute vesicular pH. Change of synaptopHluorin signal (ΔF/F) of an example KCl-responsive punctum in response to NH4Cl and MES over time is shown. These measurements were used to compute lumenal pH according to Mitchell *et al.*^62^. (**h**) Synaptic vesicle pH of wild type, *vst-1^1^* mutants, and *eat-4* mutant BAG neurons (*n* = 112, 71, and 110, respectively). Asterisk indicates *p* < 0.05 for comparisons of the mutants to wild type as determined by one-way ANOVA corrected for multiple comparisons via Dunnett’s test (the pH of all genotypes have a normal distribution, p-value for wild type vs *vst- 1^1^*: 0.041).

We considered the possibility that VST-1 regulates glutamate release by controlling the amount of vesicle fusion elicited by synaptic depolarization, *e.g.* by regulating release probability or the size of the pool of vesicles competent for fusion. To test this, we expressed the exocytosis reporter synaptopHluorin^41^ in cultured BAG neurons and asked whether *vst-1* mutation altered the amount of depolarization-evoked vesicle fusion. Wild-type and *vst-1* mutant BAG neurons displayed similar synaptopHluorin signals upon depolarization (**Supplemental Figure 3d, e**), strongly suggesting that the increase in glutamate release caused by loss of VST-1 was not the result of increased vesicle fusion. *eat-4* mutant BAG neurons showed a similar increase in synaptopHluorin signal, suggesting vesicle fusion is intact in the absence of EAT-4/VGLUT (**Supplemental Figure 3d, e**). Based on these data, we concluded that the vesicular transporter VST-1 regulates the amount of glutamate stored in synaptic vesicles, not the amount of vesicular fusion elicited by depolarization.

VGLUTs are members of a large family of anion transporters that move solutes as diverse as inorganic phosphate, acidic sugars, negatively charged amino acids, and phosphorylated adenosine nucleotides^33^. As a member of the SLC17 family of transporters, VST-1 is likely an anion transporter, and there are different ways an anion transporter in the synaptic vesicle membrane could limit glutamate uptake. VST-1 might compete with EAT-4/VGLUT for access to the electrochemical gradient used for glutamate import into synaptic vesicles. Alternatively, VST-1 might harness the same electrochemical gradient to promote glutamate efflux from synaptic vesicles. These different mechanisms will impact vesicular pH in different ways, suggesting an experimental approach to determine whether VST-1 mediates import of anions into the vesicle or instead functions as a glutamate efflux transporter. The former mechanism would increase the anion concentration in the synaptic vesicle and facilitate proton import by the vesicular V_0_-ATPase, thereby acidifying synaptic vesicles. By contrast, a glutamate efflux transporter operating as other SLC17-type anion efflux transporters, *e.g.* Sialin/SLC17A5, is predicted to have the opposite effect on vesicular pH. To distinguish these possibilities, we used synaptopHluorin to measure vesicular pH in wild-type and *vst-1* BAG neurons. Measurements of total and surface-accessible pHluorin (**Figure 3g**) allow computation of vesicular pH^42^. We found that loss of VST-1 caused a measurable increase in vesicular pH (**Figure 3h**), consistent with a model in which VST-1 supports anion influx into synaptic vesicles. We also measured vesicular pH in BAG neurons lacking EAT-4/VGLUT (**Figure 3h**). Unlike loss of VST-1, loss of EAT-4/VGLUT did not cause a measurable change in vesicular pH. These data are consistent with a model in which EAT-4/VGLUT functions with other anion transporters in the vesicle membrane and further suggest that the anion substrate for VST-1 is abundant, perhaps even more abundant than glutamate. Taken together, the effects of *vst-1* mutation on glutamate release and vesicular pH indicate that VST-1 is a vesicular transporter that competes for the electrochemical gradient required for glutamate uptake into synaptic vesicles.

### Behavioral defects of vst-1 mutants are caused by excess signaling through AMPAR-type glutamate receptors

Loss of VST-1 markedly increases evoked glutamate release from chemosensory BAG neurons. Does excess glutamate release cause the chemotaxis defects of *vst-1* mutants? We reasoned that, if so, attenuating postsynaptic glutamate receptors might restore chemotaxis to *vst-1* mutants (**Figure 4a**). The *C. elegans* genome encodes metabotropic and ionotropic glutamate receptors^20, 26, 43^. We tested our hypothesis by targeting GLR-1, an AMPAR-type receptor that is expressed by many neurons post-synaptic to BAGs^20, 44^. GLR-1 receptors also function in circuits that process inputs from glutamatergic sensory neurons^24, 45^.

**Figure 4.**
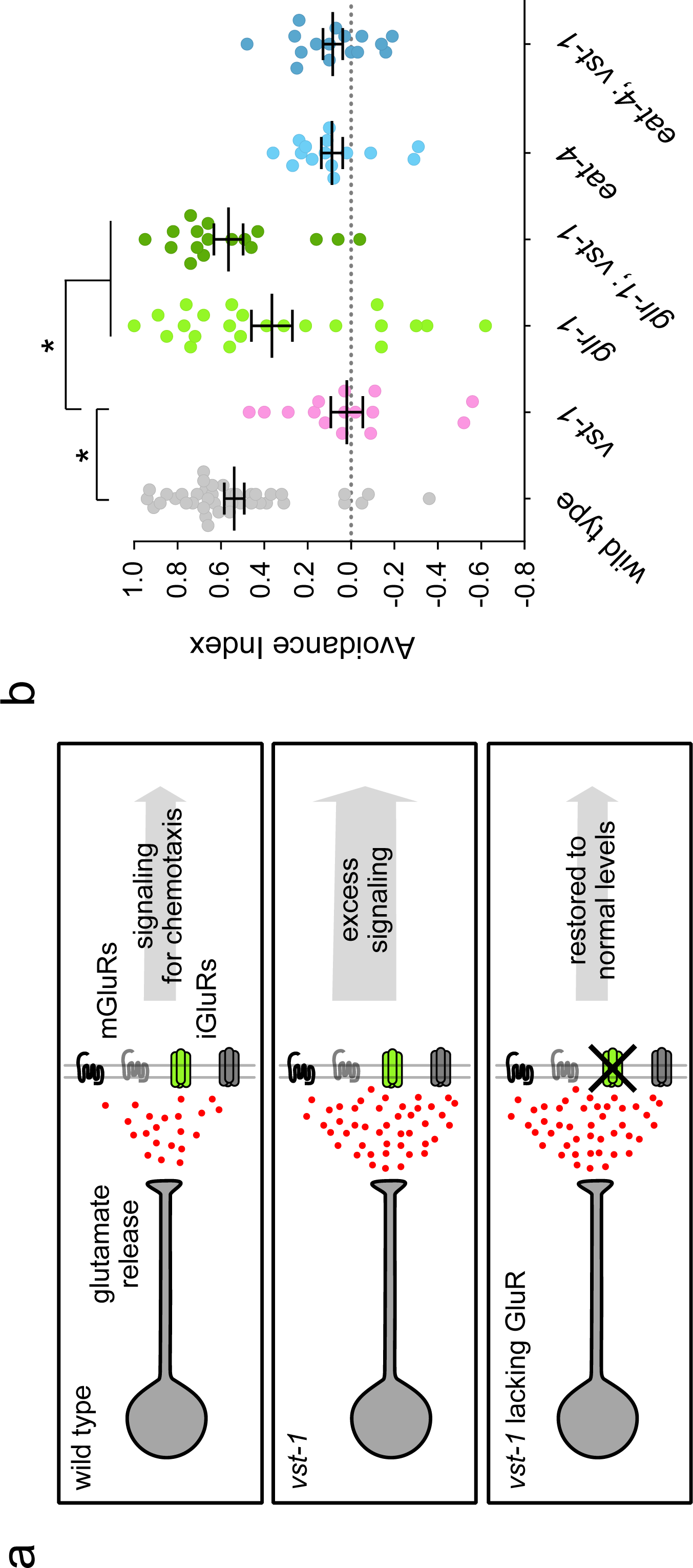
The CO_2_ chemotaxis defect of *vst-1* mutants requires an AMPAR-type glutamate receptor. (**a**) Schematic of a test for the hypothesis that excess glutamate signaling causes chemotaxis defects by over-activating glutamate receptors. (**b**) Avoidance indices (mean ± SEM) of *vst- 1(gk308047)*, *glr-1(n2461)* and *eat-4(ky5)* mutants during chemotaxis away from 10% CO_2_ (*n* = 40, 15, 23, 17, 15, and 16 for wild type, *vst-1*, *glr-1*, *glr-1;vst-1*, *eat-4*, and *eat-4;vst-1*, respectively). Asterisks indicate *p* < 0.05 from comparisons of the indicated strains to *vst-1* mutants as determined by a Kruskal-Wallis test corrected for multiple comparisons via Dunn’s test (p-values for each comparison marked with an asterisk – *vst-1^2^* vs wild type: <0.0001; *vst- 1^2^* vs *glr-1*: 0.019; *vst-1^2^* vs *glr-1;vst-1^2^*: 0.0002).

When we tested whether GLR-1 receptors were required for BAG-mediated chemotaxis, we found that *glr-1* mutants were able to avoid CO_2_ nearly as well as the wild type (**Figure 4b**), indicating that GLR-1 receptors are dispensable for the chemotaxis circuit downstream of BAGs. Strikingly, loss of GLR-1 receptors restored chemotaxis behavior to *vst-1* mutants (**Figure 4b**), consistent with a model in which loss of VST-1 causes chemotaxis defects via excess activation of glutamate receptors. By contrast, loss of EAT-4/VGLUT did not restore CO_2_-avoidance to *vst-1* mutants, confirming the essential role of glutamate signaling in BAG-mediated avoidance behavior. Together, these data strongly suggest that exuberant glutamate release by *vst-1* mutant neurons causes behavior defects *in vivo* through excess activation of AMPAR-type receptors.

To identify specific motor programs that are impacted by excess glutamate signaling in *vst-1* mutants, we used high-resolution video tracking to measure acute behavioral responses to CO_2_ stimuli of the wild type, as well as *vst-1* and *glr-1* mutants. Animals in a chamber were exposed to pulses of CO_2_-enriched atmosphere while their locomotion was recorded. From the recorded trajectories, we computed linear speed and instantaneous frequency of high-angle turns. Each of these parameters changed during a wild-type response to CO_2_-stimuli: upon sensing CO_2_, the speed of foraging animals decreased and their turn-frequency increased (**Figure 5a, d**). Slowing and turning responses required the BAG-specific CO_2_ receptor, GCY-9 (**Supplemental Figure 4**). We observed that mutation of *vst-1* affected parameters of foraging behavior prior to the CO_2_ stimulus, *i.e.* basal foraging behavior, as well as parameters of CO_2_- evoked behavior. Basal and evoked speeds of *vst-1* mutants were significantly lower than those of the wild type (**Figure 5a, d**). Also, after repeated stimulation the evoked turning-response of *vst-1* mutants was significantly greater than that of the wild type, suggesting that these mutants became partially sensitized to CO_2_ stimuli in a manner that the wild type did not (**Figure 5a, d**).

**Figure 5.**
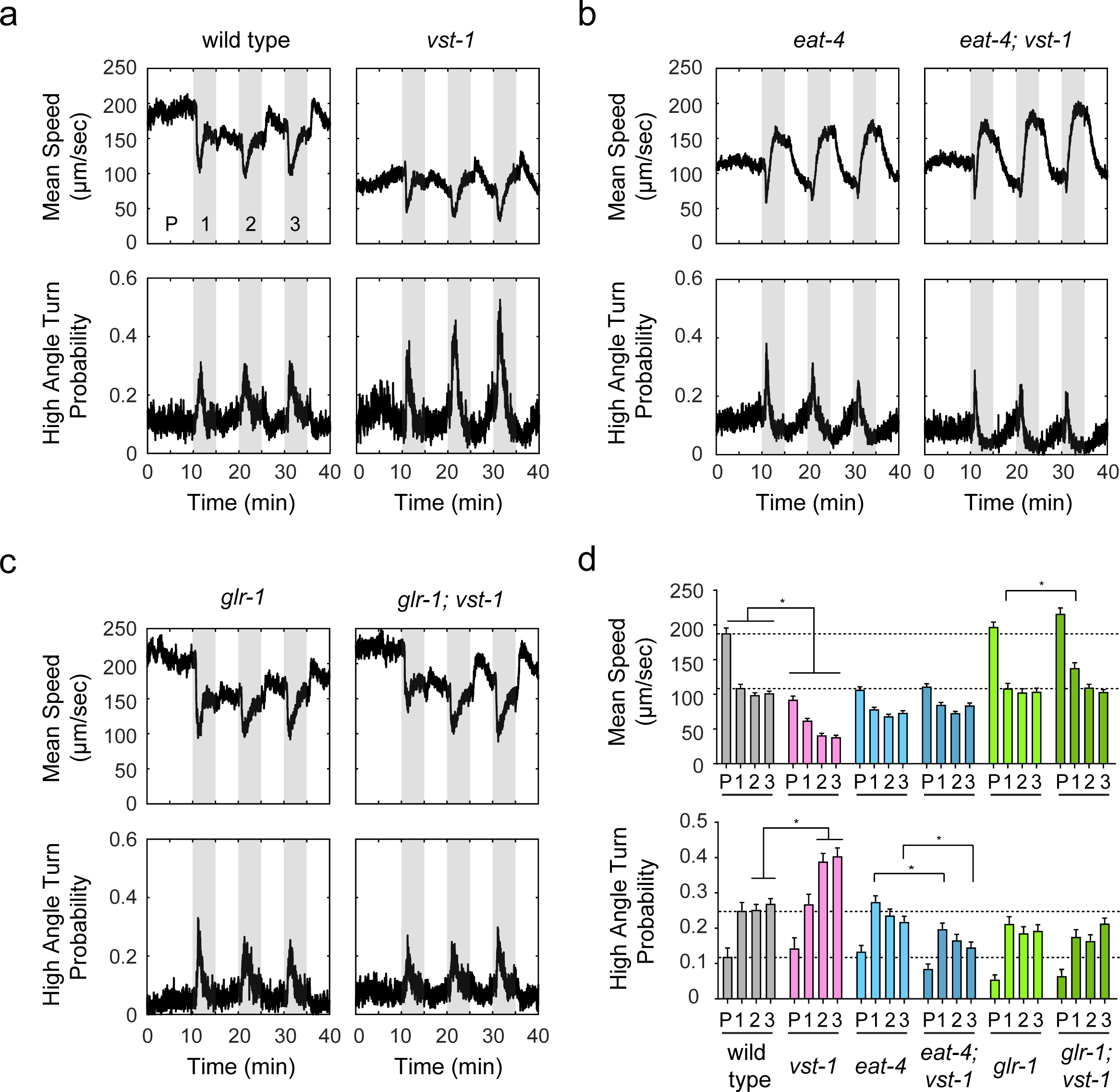
Loss of VST-1 disrupts acute behavioral responses to CO_2_ in a manner than requires GLR-1/AMPAR and EAT-4/VGLUT. (**a**) Instantaneous speed and probabilities of executing a high-angle turn of wild type and *vst-1* animals in response to three pulses of CO_2_. Pulses of CO_2_ are labeled numerically and the pre-stimulus period is labeled ’P.’ Traces in this panel and all other panels show the mean of measurements taken from at least eight independent trials of 30-60 animals. (**b**) Mean instantaneous speed and turn probabilities of *eat-4* and *eat-4; vst-1* mutants. (**c**) Mean instantaneous speed and turn probabilities of *glr-1* and *glr-1; vst-1* mutants. (**d**) Summary data from experiments shown in panels a-c. Bar graphs show mean speeds and turn probabilities ± SEM of different strains during the pre-stimulus period (P) and after each presentation of CO_2_ (labeled 1-3). The number of tracks analyzed (*n*) for mean speed during each period (P, 1, 2, and 3) are: 74, 105, 144, and 159 (wild type); 62, 79, 98, and 90 (*vst-1*); 147, 166, 151, and 163 (*eat-4*); 115, 130, 121, and 110 (*eat-4; vst-1*); 73, 85, 90, and 95 (*glr-1*); 79, 84, 99, and 131 (*glr- 1; vst-1*). The number of tracks analyzed (*n*) for high angle turn probability during each period (P, 1, 2, and 3) are: 74, 98, 145, and 158 (wild type); 62, 72, 100, and 90 (*vst-1*); 147, 166, 155, and 170 (*eat-4*); 115, 131, 120, and 107 (*eat-4; vst-1*); 73, 85, 93, and 97 (*glr-1*); 79, 84, 102, and 127 (*glr-1; vst-1*). Dashed lines indicate the means of wild type during periods P and 1. Asterisks indicate *p* < 0.05 for comparisons between genotypes during the same period determined by a Kruskal-Wallis test and corrected for multiple comparisons via Dunn’s test. Only statistical comparisons between genotypes shown in the same panel in a-c are indicated here for simplicity. Statistical comparisons between genotypes shown in different panels (*e.g.*, comparison between *vst-1* and *eat-4; vst-1*) and exact p-values for all comparisons are shown in **Supplementary Table 1**.

These data show that VST-1 is required for normal foraging behavior as well as for acute behavioral responses to CO_2_ stimuli.

Importantly, many effects of *vst-1* mutation depended upon synaptic glutamate signaling. *eat-4/VGLUT* mutants displayed abnormal slowing responses to CO_2_ stimuli and their turning responses to repeated CO_2_ stimuli also differed from those of the wild type by partially adapting (**Figure 5b, d**). Mutation of *vst-1* in *eat-4* mutants had no further effect on basal speed or on CO_2_-evoked slowing (**Figure 5b, d**). We did observe a small but significant effect of *vst-1* mutation on CO_2_-evoked turning of *eat-4* mutants (**Figure 5b, d**), but in *eat-4* mutants, loss of VST-1 caused a decrease in the frequency of turning, unlike in the wild type, whose evoked turning was greatly increased by loss of VST-1. Mutants lacking the AMPAR-type receptor GLR-1 displayed grossly normal basal and CO_2_-evoked behavior. Importantly, loss of VST-1 had no significant effect on the behavior of *glr-1* mutants (**Figure 5c, d**). These data indicate that the locomotory defects of *vst-1* mutants was suppressed by loss of GLR-1 (**Figure 5c, d**). Together, these data indicate that the chemotaxis defects of *vst-1* mutants and their defects in basal locomotion and acute responses to CO_2_ are likely caused by inappropriate activation of AMPAR-type glutamate receptors.

### Loss of VST-1 increases AMPAR-dependent synaptic glutamate signaling from BAG sensory neurons to RIA interneurons

Are there specific cells in the chemotaxis circuit downstream of BAG neurons that are affected by dysregulated glutamate signaling in *vst-1* mutants? To answer this, we measured physiological responses to CO_2_ stimuli of interneurons postsynaptic to BAGs. A subset of interneurons that receive synaptic input from BAGs^46^ express GLR-1 glutamate receptors^20, 44^, and because the *vst-1* phenotype requires GLR-1, we focused our studies on that subset of BAG targets (**Figure 6a**). We used a microfluidic device to deliver CO_2_ stimuli to immobilized animals while recording neuronal calcium responses. Under the conditions that we used, CO_2_ stimuli elicited rapid and robust responses of BAG neurons (**Supplemental Figure 5a**). During stimulus presentation, wild-type and *vst-1* mutant BAGs showed similar responses to CO_2_ stimuli (**Supplemental Figure 5a, b**), suggesting that loss of VST-1 did not grossly affect chemotransduction in BAGs. We noted that recovery of BAG calcium to baseline was slower in *vst-1* mutants, although we could not rule out the possibility that this difference was the result of experimental and sampling error (p = 0.051) (**Supplemental Figure 5c**). This effect was not observed until later in the experiment and was possibly the result of altered feedback onto BAGs in *vst-1* mutants. For this reason, we quantified interneuron activity during the time of stimulus presentation, when *vst-1* mutation had no measurable effect on BAG neuron activity.

**Figure 6.**
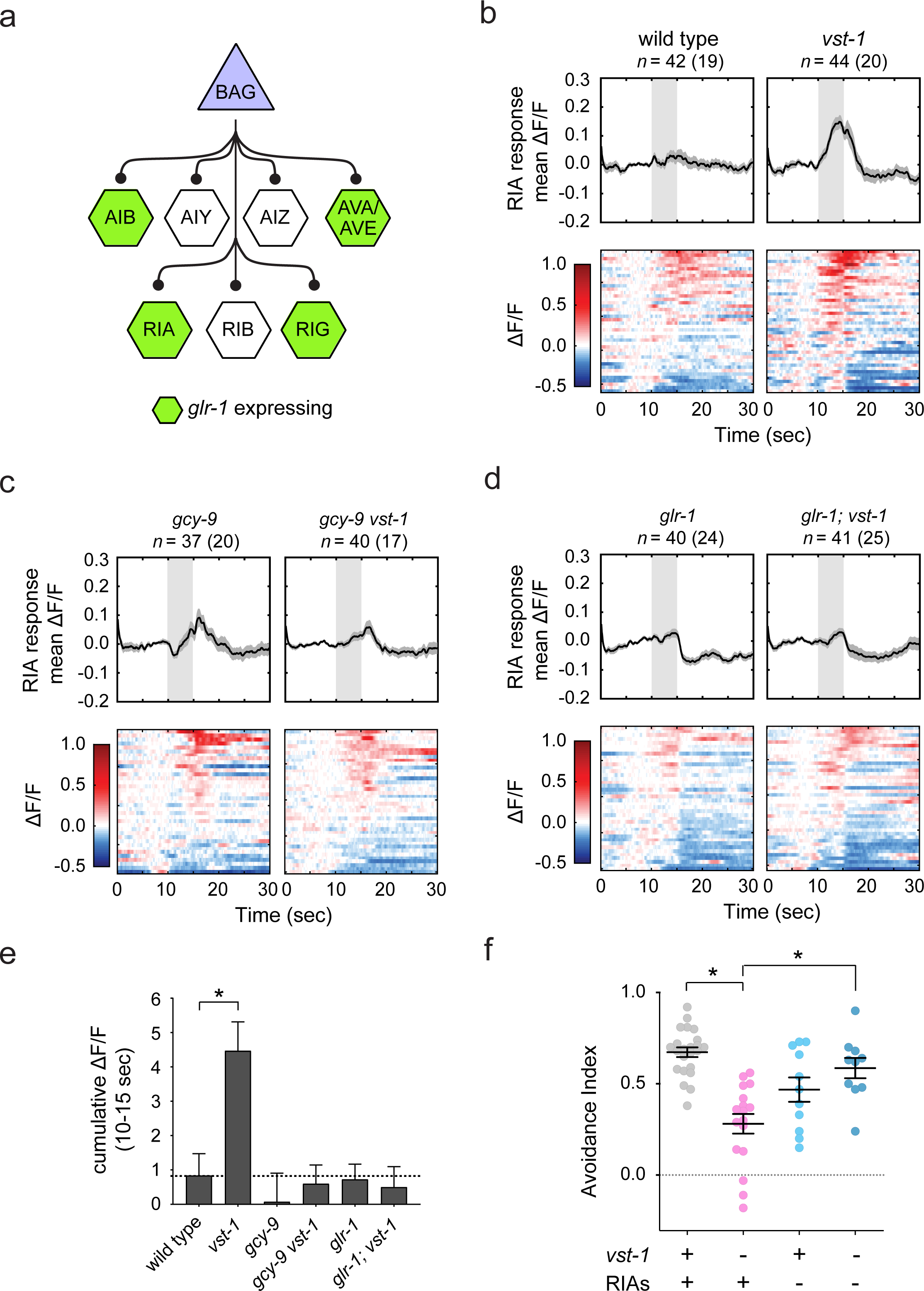
Loss of VST-1 alters the chemotaxis circuit downstream of BAGs by engaging RIA interneurons. (**a**) A schematic showing interneurons that receive synaptic input from BAG sensory neurons. (**b-d**) GCaMP signal (ΔF/F) measured in RIA neurons (*Pglr-3a::GCaMP3*) during stimulation of BAGs with 10% CO_2_ stimuli. Upper panels show the mean signal ± SEM as black lines and grey shaded regions, respectively. Individual traces were passed through a 0.5 sec moving-average filter for the plots (raw traces were used for quantification of GCaMP signal in **e**). Lower panels show individual responses as colormaps. The number in parentheses indicates the number of worms analyzed and *n* indicates the total number of cells analyzed. (**e**) Summary data from experiments shown in panels b-d. The bar graphs show cumulative GCaMP signal during CO_2_ presentation (10-15 sec) for each trial of animals of the indicated genotypes. A dashed line indicates the mean of measurements of the wild type. Asterisks indicate p < 0.05 determined by a Kruskal-Wallis test and corrected for multiple comparisons between wild type and other genotypes via Dunn’s test (p-value for wild type vs *vst-1*: 0.017). (**f**) Effects of RIA ablation on CO_2_ chemotaxis of the wild type and *vst-1* mutants. The mean ± SEM of avoidance indices for each condition are indicated (*n* = 23, 17, 11, and 10 for wild type, *vst-1*, RIA ablation in wild type background, and RIA ablation in *vst-1* mutants, respectively). Asterisks indicate p < 0.05 determined by a Kruskal-Wallis test and corrected for multiple comparisons between *vst-1* mutants and other genotypes via Dunn’s test (p-values for each comparison marked with an asterisk – *vst-1* vs wild type: <0.0001; *vst-1* vs *vst-1;* RIA_-_: 0.014).

Several interneuron-types displayed responses to BAG activation that were unaffected or only modestly affected by *vst-1* mutation. Loss of VST-1 did not significantly affect the variable responses of AIB interneurons to BAG activation (**Supplemental Figure 6a**). Calcium levels in AVA interneurons frequently dropped after BAG activation (**Supplemental Figure 6b**), suggesting that these interneurons receive inhibition from BAGs. AVA responses to BAG activation, like AIB responses, were not strongly impacted by loss of VST-1. AIY interneurons were stimulated by BAGs and displayed more consistent increases in cell calcium in response to BAG activation. AIY responses were, on average, slightly increased by loss of VST-1, but we could not rule out the possibility that the observed differences resulted from experimental and sampling errors (**Supplemental Figure 6c**).

By contrast, we observed a clear effect of *vst-1* mutation on signaling from BAGs to RIA interneurons. In the wild type, RIAs responded variably to BAG activation with either increases or decreases in cell calcium; on average, these responses were balanced and the mean response of RIAs to BAG activation was close to zero (**Figure 6b, e**). Loss of VST-1 caused a significant increase in excitation received from BAGs, and the mean response of RIAs in *vst-1* mutants indicated the appearance of more prominent excitatory input to RIAs (**Figure 6b, e**). To demonstrate that this effect of *vst-1* mutation was the result of increased signaling from BAGs, we measured RIA responses to CO_2_ stimuli in *gcy-9* mutants, which lack a key component of the CO_2_-sensing machinery in BAGs^36, 37^. Loss of GCY-9 in *vst-1* mutants suppressed the effect of *vst-1* mutation, suggesting the BAG CO_2_-sensing machinery is required for its effect (**Figure 6c, e**). We further tested whether the effects of *vst-1* mutation on RIA activation by BAGs require GLR-1 glutamate receptors, as predicted by our model. In mutants lacking GLR-1, there was no clear effect of *vst-1* mutation (**Figure 6d, e**), indicating that the increased activation of RIAs observed in *vst-1* mutants requires signaling through GLR-1.

To determine whether increased activation of RIA contributes to the chemotaxis defects of *vst-1* mutants, we tested the effect of *vst-1* mutation on CO_2_ avoidance by the wild type and by animals lacking RIA interneurons. As we observed previously, *vst-1* mutants displayed a robust chemotaxis defect (**Figure 6f**). Animals lacking RIAs in a wild-type background did not display a clear defect in chemotaxis behavior (**Figure 6f**). Notably, loss of RIAs in *vst-1* mutants restored CO_2_ chemotaxis (**Figure 6f**). These data indicate that the chemotaxis defect of *vst-1* mutants is caused, at least in part, by ectopic activation of RIA interneurons.

## DISCUSSION

### A presynaptic mechanism sets the strength of glutamatergic synapses via regulation of neurotransmitter loading

Proper regulation of synaptic strength is critical for the function of neural circuits, and mechanisms that change synaptic strength are drivers of circuit plasticity and are required for adaptation and learning. Well understood plasticity mechanisms remodel or modulate the complement of postsynaptic neurotransmitter receptors to alter how synaptic signals are processed by the receiving neuron. Such mechanisms play critical roles in glutamatergic synapses, and their molecular underpinnings have been extensively studied (reviewed in ^47^). Other plasticity mechanisms regulate the kinetics of synaptic vesicle fusion, either by altering presynaptic calcium levels or by modulating the fusion machinery itself (reviewed in ^48^). Our study of a chemotaxis circuit in *C. elegans* suggests that the strength of glutamatergic synapses can also be determined by a mechanism that sets the amount of glutamate packaged into synaptic vesicles. We find that the amount of glutamate released by chemosensory neurons is controlled by interactions between VGLUT and another vesicular transporter - VST-1 - that opposes VGLUT function. Increasing glutamatergic neurotransmission by disrupting the balanced antagonism between VGLUT and VST-1 causes a behavioral defect that was as severe as that caused by mutations that eliminate glutamatergic neurotransmission, *e.g.* VGLUT mutation. This observation demonstrates the importance of mechanisms that assign appropriate strengths to functional connections within the chemotaxis circuit downstream of BAG neurons.

How does VST-1 oppose the function of VGLUT to determine the strength of glutamatergic signaling? Functional interactions between these transporters must be considered in the context of what is known about the energetics of neurotransmitter transport. It is possible that VST-1 antagonizes VGLUT by driving glutamate export from vesicles and directly counteracting VGLUT-mediated glutamate import. This ’revolving-door’ model for glutamate transport across synaptic vesicle membranes demands that VST-1 be a glutamate efflux transporter. We do not favor this model for two reasons. First, it seems highly unlikely that VST- 1 is a glutamate transporter. Recent structural studies of VGLUT have identified key residues implicated in glutamate binding^49^, and these residues are not conserved between VST-1 and VGLUT (**Supplemental Figure 7**). Second, our analysis of the effects of VST-1 mutation on vesicular pH indicated that loss of VST-1 alkalinizes vesicles, which is consistent with VST-1 mediating anion influx, not glutamate efflux. We therefore favor a model in which VST-1 competes with VGLUT for access to Δ Ψ, the electrical potential gradient across the vesicular membrane (**Figure 7**). An important implication of this model is that it predicts that the abundance of VST-1 substrates would regulate the glutamate content of synaptic vesicles and synaptic transmission in a chemotaxis circuit. Such substrates would, therefore, function as endogenous small-molecule regulators of glutamatergic neurotransmission.

**Figure 7.**
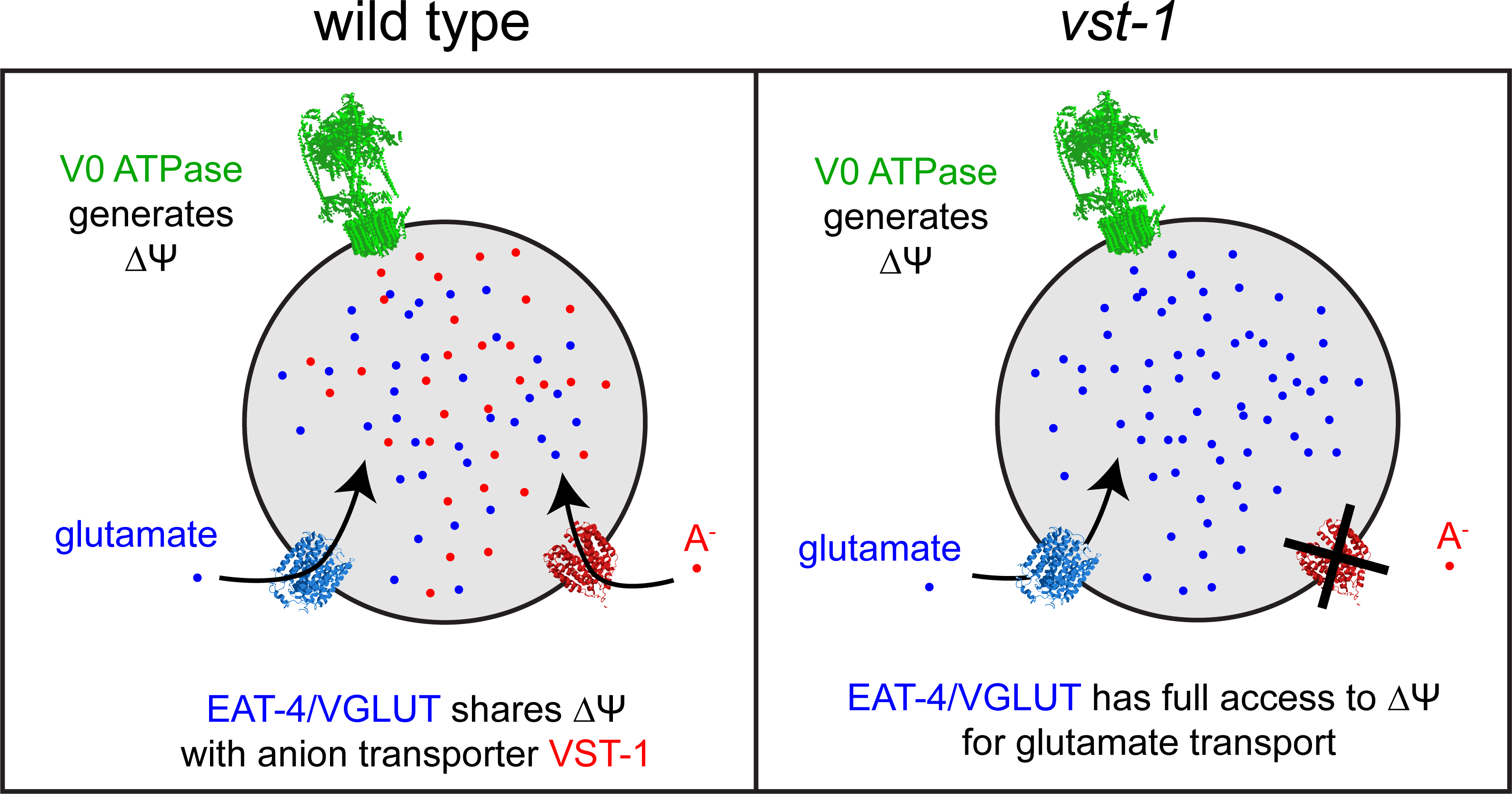
A model for VST-1 regulation of glutamate transport into synaptic vesicles. A transmembrane potential Δ Ψ generated by the vesicular ATPase is harnessed by VGLUT to transport glutamate (illustrated as blue dots) into synaptic vesicles. In our model VST-1- mediated transport of an anion (red dots) into the vesicle dissipates Δ Ψ and thereby reduces glutamate transport.

### The strength of synaptic glutamate signaling is tuned to support BAG-dependent chemotaxis

Loss of VST-1 causes a chemotaxis defect that is as severe as that caused by loss of VGLUT. This observation indicates that the strengths of glutamatergic synapses in the chemotaxis circuitry must be set within a certain range in order to correctly process inputs from BAG sensory neurons. BAG-dependent chemotaxis, like other chemotaxis behaviors^50–52^, is the result of a sensory system coordinating changes to motor programs that control speed and turning. We found that decreased and increased glutamate signaling caused by loss of VGLUT and loss of VST-1, respectively, did not eliminate the behavioral components of chemotaxis but instead disrupted their coordination. Loss of glutamate signaling had a dramatic effect on speed-control and resulted in animals that were incapable of sustained slowing during BAG stimulation. By contrast, increased glutamate signaling caused by *vst-1* mutation resulted in persistent and stimulus-independent slowing. *vst-1* mutants also displayed exaggerated turning responses upon BAG stimulation, an effect that was not anticipated by our measurements of turning by *eat-4/VGLUT* mutants. It is noteworthy that the behavior defects caused by loss of EAT- 4/VGLUT and VST-1 are not precise mirror images of each other. This might indicate that loss of VST-1 does not affect all glutamatergic synapses equally, unlike loss of EAT-4/VGLUT, which is required across the board for all types of glutamatergic neurotransmission. Importantly, our data indicate that the effects of *vst-1* mutation on behavior require the AMPAR-type glutamate receptor GLR-1 and are occluded by the effects of *eat-4/VGLUT* mutation, strongly indicating that VST-1 dysregulates synaptic glutamate signaling *in vivo*.

Our analysis of neurons that receive synaptic inputs from BAGs might also suggest that synapses in the chemotaxis circuit are differently sensitive to loss of VST-1. We found that BAG signaling to one set of targets - the RIA interneurons - was significantly affected by loss of VST-1, while signaling from BAGs to other targets was either not affected or only affected slightly (**Supplemental Figure 6**). In *vst-1* mutants, RIAs on average received more excitatory input from BAGs (**Figure 6b**), and this excitation required the AMPAR-type glutamate receptor GLR- 1 (**Figure 6d**), which is a critical mediator of fast excitatory signaling in the *C. elegans* nervous system^22–24^. These observations might suggest that VST-1 has a privileged role at certain excitatory synapses in the chemotaxis circuit.

### Interactions between vesicular transporters as a mechanism for regulating neurotransmission

We suggest that the functional interaction between VGLUT and VST-1 in glutamatergic synapses is an example of a more general mechanism that regulates neurotransmission by controlling neurotransmitter packing into synaptic vesicles. There is evidence for such a mechanism from prior studies of monoaminergic and cholinergic neurotransmission. Dopaminergic neurons in the vertebrate and insect brain co-express VMAT and VGLUT^6, 7^. Eliminating VGLUT from dopaminergic neurons reduces dopamine storage in presynaptic terminals and diminishes dopamine release. VGLUT facilitates dopamine storage by facilitating the energetics of VMAT-dependent dopamine transport. VMAT relies on a proton gradient to transport dopamine into vesicles, and vesicular glutamate offsets the charge imbalance caused by proton transport permitting a steeper proton gradient and, as a consequence, more dopamine transport. A similar interaction between VGLUT and VAChT has been proposed to explain how VGLUT boosts cholinergic signaling in the vertebrate striatum^5^. VGLUT-VMAT and VGLUT- VAChT interactions illustrate how vesicular transporters that access different components of the electrochemical gradient that powers neurotransmitter uptake can mutually reinforce their distinct transport functions.

Besides these examples, how widespread are functional interactions between synaptic vesicular transporters? This is a difficult question to answer, in no small part because the full complement of synaptic vesicle-associated transporters remains to be determined. There is no sequence-based hallmark of vesicular transporters, and these factors are expressed at low levels, which confounds biochemical approaches for their identification. New vesicular transporters continue to be discovered; in addition to VST-1, recent studies of neural circuits in the insect brain have identified a transporter named Portabella^12^, which is related to VMAT and VAChT, and a transporter named LOVIT, which is expressed in histaminergic neurons^13^. The continued characterization of the large and diverse family of transporter proteins will likely add to the census of transporters in synaptic vesicle membranes, and each of these transporters is in principle capable of modulating neurotransmission via the mechanisms discussed.

With respect to VST-1, our data suggest that this particular transporter has functions in non-glutamatergic neurons in addition to its function in glutamatergic neurons. VST-1 is clearly expressed by many non-glutamatergic neurons (**Figure 2a**), raising the possibility that VST-1, like vertebrate VGLUT isoforms, might potentiate neurotransmitter packaging mechanisms that require proton gradients, *e.g.* monoamine and acetylcholine transport. Therefore, depending on the energetics of the neurotransmitter transporters in a synaptic vesicle membrane, the function of VST-1 might change from antagonizing neurotransmitter uptake, as it does glutamate uptake, to potentiating it. In a *C. elegans* chemotaxis circuit, VST-1 is required as an auxiliary that regulates VGLUT-dependent glutamate signaling. Such auxiliary transporters might be widespread regulators of synaptic neurotransmission in other circuits, and it is intriguing to consider the possibility that their functional interactions with neurotransmitter transporters constitute mechanisms that generally regulate synaptic strength *in vivo*.

## METHODS

### C. elegans strains

Mutant and transgenic animals used for this study are listed in **Supplementary Table 2**. Transgenic animals used for experiments were generated by microinjection following the standard protocol. For most lines generated by injection of a plasmid, the plasmid was injected at 20-40 ng/μl concentration. The exceptions were:

RNAi plasmids (pJC141, pJC143, pJC133, pJC135) – 100 ng/μl

pJC151 (*Pgcy-33::snb-1::superecliptic pHluorin*) - 5 ng/μl

*Punc-122::GFP, Punc-122::mCherry* - 60 ng/μl

For the *vst-1* fosmid reporter line (FQ1571), a linearized *vst-1* fosmid reporter (pEH62) was injected at 11 ng/μl concentration into a *eat-4* fosmid reporter line (OH11124; *otIs388[eat-4 fosmid::SL2::YFP::H2B + (pBX) pha-1(+)] pha-1(e2123)*) with carrier DNA (*vst-1(gk308047)* genomic DNA digested with EcoRV, injected at 202 ng/μl) and co-injection plasmid (*Punc- 122::GFP*, injected at 65 ng/μl). For lines with the *vst-1::GFP* translational reporter (FQ1137, FQ1149, FQ1167), a fosmid (pJC77) carrying the reporter was amplified by PCR using primers pJC206 and pJC207 to obtain a 9.6 kb product (pJC106). For lines with the *eat-4::mCherry* translational reporter (FQ1167), a fosmid (pJC52) carrying the reporter was amplified by PCR using primers pJC221 and pJC220 to obtain a 14.2 kb product (pJC149). These PCR products were gel-purified and injected at 54 (pJC106) and 44 (pJC149) ng/μl concentrations, respectively, to generate each line.

### Plasmids and primers

The plasmids, fosmids, and primers used for this study are listed in **Supplementary Table 3**. The sense and antisense plasmids used for RNAi microinjection were constructed according to Esposito *et al.*^53^. *vst-1* RNAi plasmids, pJC141 and pJC143, were generated by amplifying *vst-1* cDNA by PCR using JC178/JC179 (sense) and JC180/JC181 (antisense) primer sets, respectively, and targeted 356 nucleotides spanning exons 2 to 4 of *vst-1* (nucleotides 103 to 458 of cDNA). *eat-4* RNAi plasmids, pJC133 and pJC135, were generated by amplifying *eat-4* cDNA by PCR using JC170/JC171 (sense) and JC172/JC173 (antisense) primer sets, respectively, and targeted 350 nucleotides spanning exons 3 to 5 of *eat-4* (nucleotides 211 to 560 of cDNA). The amplified sense or antisense fragments were placed behind a *flp-17* promoter by Gibson assembly^54^. GFP or mCherry cassettes were inserted into the *vst-1* and *eat-4* fosmids according to Tursun *et al.*^55^.

### RNASeq data analysis

From the RNASeq dataset in Horowitz *et al.*, 2019^32^ (GEO accession: GSE137267; Accession for BAG dataset: GSM4074164 and GSM4074165; Accession for unsorted cells dataset: GSM4074162 and GSM4074163), data analysis was performed essentially as described previously^56^. The coverage histograms for *eat-4* and *vst-1* were visualized using the Integrative Genomics Viewer^57^ and the mean read counts were quantified using Deeptools^58^. Fold enrichment and false discovery rate were computed using the DESeq2 package^59^.

### Phylogenetic tree

The phylogenetic tree in **Figure 1d** was generated using the multiple sequence alignment program T-Coffee (www.ebi.ac.uk/Tools/msa/tcoffee).

### CO_2_ chemotaxis assays

20-60 adult hermaphrodites were washed in M9 solution and placed on unseeded 10 cm NGM plates and a custom-made chamber with a thin layer of glycerol applied to the edges to prevent animals from escaping was pressed into the NGM agar plates. The chamber was 6 cm in diameter and had two gas inlets 2 cm from the center and on opposite sides. Air (20% O_2_, N_2_ balance) flowed into one inlet and CO_2_ (10% CO_2_, 20% O_2_, N_2_ balance) flowed into the other at 1.5 ml/min using a dual syringe pump (New Era). The number of worms on each side were counted after 30 min and an avoidance index (AI) was computed by:

AI = (number of worms on air side – number of worms on CO_2_ side) / (total number of worms). Kruskal-Wallis and Dunn’s multiple comparison test were used for statistical comparisons between wild type (or *vst-1* depending on the experiment) and other genotypes. A few trials were represented in multiple figures because multiple experiments were simultaneously conducted on a single day: one trial for wild type and two trials for *vst-1* were used for **Figures 1e, 4b, and 6f**, and two trials each of wild type, *vst-1*, and *eat-4* were used for both **Figures 1e** and **4b**. RIA ablation was confirmed before conducting chemotaxis assays for the ablation lines by checking for mCherry expression in RIAs under a fluorescent dissecting microscope (high power).

### Quantification of cellular expression of VST-1 in glutamatergic neurons

Coexpression of *vst-1* and *eat-4* nuclear reporters was measured by manually labeling each YFP-positive, mCherry-positive, and double positive nucleus with a unique ID (visible in both YFP and mCherry channels and on all optical slices) in ImageJ^60^ while progressing through the entire head one optical slice at a time.

### Electron microscopy

*Vst-1::GFP; eat-4::mCherry; Punc-122::GFP* transgenic worms were subject to high pressure freezing, freeze substitution, embedment in Lowicryl HM20, and postembedding immunolabeling according to Hall *et al.*, 2012^61^. 4-5 worms were aligned in one direction into a hat filled with yeast. The hat was frozen under high pressure (2100 bar) and subsequently immersed in liquid nitrogen. Samples were fixed with 0.2% glutaraldehyde/ 98% acetone/ 2% water at -90°C for 110 hrs, slowly warmed up to -20°C (5 °C/hr), held at -20°C for 16 hrs, slowly warmed up to 0°C (6 °C/hr), and held at 0°C. The samples were rinsed in 100% acetone 4x15min at 0°C, removed from the hat, placed in HM20 acetone (1:2 ratio), and held at 4°C for 3 hrs. Solution was changed to HM20 acetone (2:1) at 4°C for 10-20 hrs, then changed to 100% HM20 5 times over 48 hrs and held at -20°C. Samples were transferred to gelatin capsules, filled with HM20, sealed, and cured under UV light for 1-2 days at -20°C. Ultrathin sections (70 nm) were cut by a Leica UC6/FCS microtome. Immunostaining was conducted using an anti-GFP primary antibody (Millipore Sigma, ab3080, 1:10 dilution) and 15nm Protein A gold-conjugated secondary antibody (PA15, Cell Microscopy Center, University Medical Center Utrecht, 35584 CX Utrecht, The Netherlands) on a carbon-formvar coated copper grid. No primary antibody solution was used for control staining. Images were acquired using a Philips CM-12 (FEI, Eindhoven, The Netherlands) transmission electron microscope captured with a CCD camera (Gatan 4k x 2.7k digital camera, Gatan Inc., Pleasanton, CA).

### Cellular glutamate release assay

*C. elegans* embryonic cells from the *Pflp-17::iGluSnFR* line were cultured as previously described^37^ to obtain BAG neurons expressing iGluSnFR on the cell membrane directed toward the extracellular solution. Immediately prior to an assay, cells were washed ten times with 3 ml control solution to rinse out residual glutamate from the culture medium. A multichannel perfusion system (Automate Scientific) was used to deliver control (145 mM NaCl, 5 mM KCl, 2 mM CaCl_2_, 1 mM MgCl_2_, 10 mM HEPES, 10 mM glucose, pH 7.2, 335-345 mOsm) and 100 mM KCl (50 mM NaCl, 100 mM KCl, 2 mM CaCl_2_, 1 mM MgCl_2_,10 mM HEPES, 10 mM glucose, pH 7.2, 335-345 mOsm) solutions to cells on peanut lectin-coated MatTek dishes. Images were acquired every 100ms with a 20x Nikon objective (air, NA 0.8) on a custom-built microscope with a 488 nm excitation light. Excitation and image acquisition were controlled by Live Acquisition software (Till Photonics). Region-of-interest (ROI) was selected as 3x3 pixel square located at tip of neurite and mean pixel intensity was used for analysis. A custom Matlab script was used to generate a plot showing ΔF/F over time and calculate area-under-the-curve (AUC) between 10 and 20 seconds.

### pH measurements via synaptopHluorin

Dissociated BAG neurons expressing synaptopHluorin were prepared and changes in synaptopHluorin fluorescence in response to KCl, NH_4_Cl, and MES treatment were acquired as described above for the cellular glutamate release assay with the following differences. Cell cultures were obtained from *Pgcy-33::snb-1::superecliptic pHluorin; Pflp-17::mStrawberry; Punc-122::mCherry* lines. Cells were not subject to extensive washing to remove residual glutamate prior to the assay. The cell marker mStrawberry was used to identify BAG cell neurites and used to guide search for synaptopHluorin puncta within BAG neurites. 3x3 pixel regions expressing synaptopHluorin within BAG neurites were selected for analysis. Multiple puncta were selected from a single neuron if they were not adjacent.

In order to identify KCl-responsive puncta, we first assessed the response to 100 mM KCl for 5 seconds. Punctum with baseline fluorescence values close to background levels [mean (0- 10sec) ≤ mean + 3SD of background signal (0-10 sec)] were excluded because ΔF/F were too noisy and fluctuating. KCl-responsive puncta were chosen by determining whether the maximum ΔF/F between 10 and 20 seconds (after passing through a 1 sec-moving-average filter) was greater than the baseline ΔF/F + 3SD [max (10-20 sec) > mean + 3SD of baseline signal (0-10 sec) using traces passed through a 1 sec-moving average filter].

KCl-responsive puncta were further subject to pH measurements as previously described^42, 62^ with the following details. NH_4_Cl (95 mM NaCl, 50 mM NH_4_Cl, 5 mM KCl, 2 mM CaCl_2_, 1 mM MgCl_2_, 10 mM HEPES, 10 mM glucose, pH 7.2, 335-345 mOsm) was applied for 40 seconds (control saline 20 sec – NH_4_Cl 40 sec – control saline 20 sec). MES (145 mM NaCl, 5 mM KCl, 2 mM CaCl_2_, 1 mM MgCl_2_, 10 mM MES, 10 mM glucose, pH 5.5, 335-345 mOsm) was applied for 60 seconds (control saline 20 sec – MES 60 sec – control saline 120 sec). Cells were subject to additional washes by control saline (145mM NaCl, 5mM KCl, 2mM CaCl_2_, 1mM MgCl_2_, 10mM HEPES, 10mM glucose, pH 7.2, 335-345 mOsm) between different treatments to ensure clearing of the previous treatment from the perfusion system.

Fractional increase of synaptopHluorin signal in response to NH_4_Cl (γ) for each punctum was acquired by determining the maximum ΔF/F during NH_4_Cl application (20-60 sec) after passing the trace through a 5 sec-moving-average filter to remove high frequency fluctuations hindering accurate measurements. Exclusion criteria used for KCl responders were applied to this experiment to define NH_4_Cl responders: puncta with baseline fluorescence values close to background levels (within 3SD) or with a maximum ΔF/F equal or less than 3SD above baseline (after passing through a 1 sec-moving-average filter) were excluded from pH calculation. Fractional decrease of signal in response to MES (ε) was acquired by determining the minimum ΔF/F during MES application (20-80 sec) after passing the trace through the same moving-average filter and applying the same exclusion criteria as for NH_4_Cl to define MES responders. The pH for each punctum was obtained from measurements of γ and ε using equations described in Mitchell *et al.* 2004^62^. Finally, pH values calculated to be below 5.0 were excluded as they were outside the dynamic range of superecliptic pHluorin^41^. The number of KCl-responsive puncta excluded for pH measurements by these criteria were 6 (of 118), 5 (of 76), and 5 (of 115) for wild type, *vst-1*, and *eat-4*, respectively.

### Videotracking and analysis of acute locomotor response to CO_2_

Locomotory response to CO_2_ was acquired and measured essentially as previously described^63^. A custom-built worm tracker was illuminated with red ring lights and the movement of multiple worms was recorded by a CCD camera (Unibrain, Fire-I 785b). 30-60 fed adult hermaphrodites were washed with M9 solution, placed on an unseeded 10 cm NGM plate, and allowed to move within a 6 cm diameter area encircled by a thin layer of glycerol for 5-10 min so that the solution could evaporate. Within 15 min of washing worms, a 6 cm diameter chamber with a gas inlet 1 cm from its edge is pressed onto the NGM to form a seal and alternating flows of air (20% O_2_, N_2_ balance) and CO_2_ (10% CO_2_, 20% O_2_, N_2_ balance) are allowed to enter the chamber at 1.5 ml/min. Videorecording of the movement and engagement of shuttle valve (Neptune Research, SH360T041, Caldwell NJ) are controlled by a Matlab script (Nihkil Bhatla). Field of view for tracking was set to capture movement of worms in the half of the chamber closer to the gas inlet.

After the experiment, a Matlab script (Nihkil Bhatla) is used to identify worms by morphological features and calculate locomotory parameters such as instantaneous speed and heading/angle change. In order to exclude non-worm artifacts, we limited our analysis to objects that lasted at least 30 sec and had a minimum speed of 1.17 pixels/sec (38.15 μm/sec). Upon manual inspection of 200 tracks for each genotype, we found these criteria selectively exclude non-worm objects - less than 5% of excluded objects are worms for all genotypes except for *eat- 4* (7%) and *gcy-9* (5.5%). For plots showing change of speed over time, we calculated the mean instantaneous speed of the population of worms at each time point. We defined a high angle turn as a change of direction equal or greater than 50°/sec^50^ and calculated the probability that the population of worms at each time point will execute a high angle turn to generate plots showing turn probability over time. For statistical comparisons, we found the time point (t_min_ for speed and t_max_ for turn probability) at which mean instantaneous speed is minimum (or turn probability is maximum) for each stimulus window, identified tracks spanning t_min_-15 to t_min_+17 sec (or t_max_-15 to t_max_+17 sec), calculated the mean instantaneous speed (or turn probability) over that time window for each individual track, and conducted Kruskal-Wallis (and Dunn’s multiple comparisons) test among genotypes. For the pre-stimulus (P) window, tracks spanning 465-497 sec were used for quantification.

### GCaMP imaging via microfluidic device

We used microfluidic devices (MicroKosmos and custom-built) described in Chronis *et al.* 2007^64^, but modified the solution flow system to be driven by positive pressure similar to the pressure system used by Rouse *et al.*^65^ to reduce formation of bubbles in the tubing or channels. A custom-built pressure/valve control box was used to deliver filtered air into solution reservoirs which were connected to the microfluidic device via tubing. The pressure on all channels was 2- 5 psi and the pressure in the side channels (channel 1 and 4 in Chronis *et al.* 2007) were set 0.5-1 psi lower than that of the control and stimulus solution channels (channel 2 and 3) to allow rapid transition (< 300ms) between control and stimulus solutions (**Supplemental Figure 8**). Transition between solutions was mediated by a two-way pinch valve (Cole Parmer, GH-98302- 02). Stimulus solution was 18.45mM NaHCO_3_ in S Basal medium (equilibrates with 10% CO_2_ and pH 7.2 assuming closed system) and fluorescein (Cole Parmer, #00298-17, 1:10,000,000 dilution) was added to visualize presence of stimulus. Red food dye (McCormick, 930651) was used instead of fluorescein for GCaMP experiments for AVA. NaCl was added to S Basal medium and pH was adjusted to match the osmolarity and pH of the control solution to that of the stimulus solution. Images were acquired on the inverted microscope as described above using a 20x objective (Nikon, air, 0.8NA) and the illumination (488nm), image acquisition, and transition between solutions was driven by the Live Acquisition software. We performed up to three experiments per worm with at least 1 min between the stimulus presentations to allow GCaMP signals to return to baseline levels. We identified AVA neurons in the *Popt-3::GCaMP6* line based on its anterior-most location. All other lines used for GCaMP imaging were specific for the indicated neuron in **Figure 6, Supplemental Figure 5, and Supplemental Figure 6**.

Image registration was conducted using the ImageJ StackReg Plugin (‘translation’ algorithm) and ROIs were selected around the soma for BAG neurons and around the neurites for interneurons. The background ROI used for background subtraction was selected outside of the worm instead of within the worm due to the severe fluctuation of fluorescence intensities of the latter. We found the maximum ΔF/F of the former is roughly half that of the latter and acknowledge the former method does not control for the basal levels of fluorescence in the worm not associated with GCaMP expression (such as autofluorescence), but decided to use it to limit fluctuation of ΔF/F due to fluctuating levels of background fluorescence. The mean intensity of fluorescence of the ROIs were used to generate plots showing ΔF/F changes over time using a Matlab script. Occasionally, we observed huge increases of fluorescence in the absence of stimulation, perhaps correlated with head or body movement, during the recordings. In order to avoid the influence of this CO_2_-independent activity, we excluded individual traces which had big increases in ΔF/F during the pre-stimulus period [max (0-10 sec) > mode + 5SD (0-10 sec) using traces passed through a 0.5 sec-moving average filter]. The mode of the individual recording was used (instead of the mean) as baseline and the mean SD of all recordings (0-10 sec) from a genotype was used (instead of the SD of the individual recording between 0-10 sec) because a recording with a big increase in fluorescence during the pre-stimulus has a big mean and a big SD. The number of recordings excluded by this criterion for RIA were 2 (of 44), 2 (of 46), 6 (of 43), 3 (of 43), 4 (of 44), and 7 (of 48) for wild type, *vst-1*, *gcy-9*, *gcy-9 vst-1*, *glr-1*, and *glr-1; vst-1*, respectively. To better represent the variability among the individual responses, we used colormaps ranked in descending order according to the mean ΔF/F value between 10-15sec using a Matlab script.

## ACKNOWLEDGEMENTS

This work was supported by NIGMS grant R35 GM122573 and F31 NS100360 (L.B.H.). Chris Petzold, Kristen Dancel-Manning, and Alice Liang at NYULH Microscopy Laboratory, which is partially supported by Laura and Isaac Perlmutter Cancer Center Support Grant NIH/NCI P30CA016087, prepared samples for electron microscopy and acquired electron micrographs. Katie Eyring generated the *Popt-3::GCaMP6f* strain and Elver Ho generated the pEH62 fosmid. Nikhil Bhatla and Alice Fok helped us establish protocols for video tracking. We thank Loren Looger, Jeremy Dittman, Andrew Gordus, Sharad Ramanathan, and Daniel Colon-Ramos for providing plasmids and strains used in this study. We also thank Hang Lu for instruction and advice on microfluidics, Jeremy Dittman for his expertise on pHluorin analysis, and Da-Neng Wang, David Sauer, Nicolas Tritsch, Villu Maricq, and Oliver Hobert for critical discussions. Some strains used in this study were provided by the *Caenorhabditis* Genetics Center, which is supported by the NIH Office of Research Infrastructure Programs grant P40 OD010440.

## AUTHOR CONTRIBUTIONS

J.-H. C. designed and performed all experiments and analyses except BAG transcriptome studies, which were performed by L.B.H.; N.R. contributed to experimental design and data analysis. J.-H. C. and N. R. formatted data for presentation and wrote the manuscript.

## SUPPLEMENTAL FIGURE LEGENDS

**Supplemental Figure 1.**
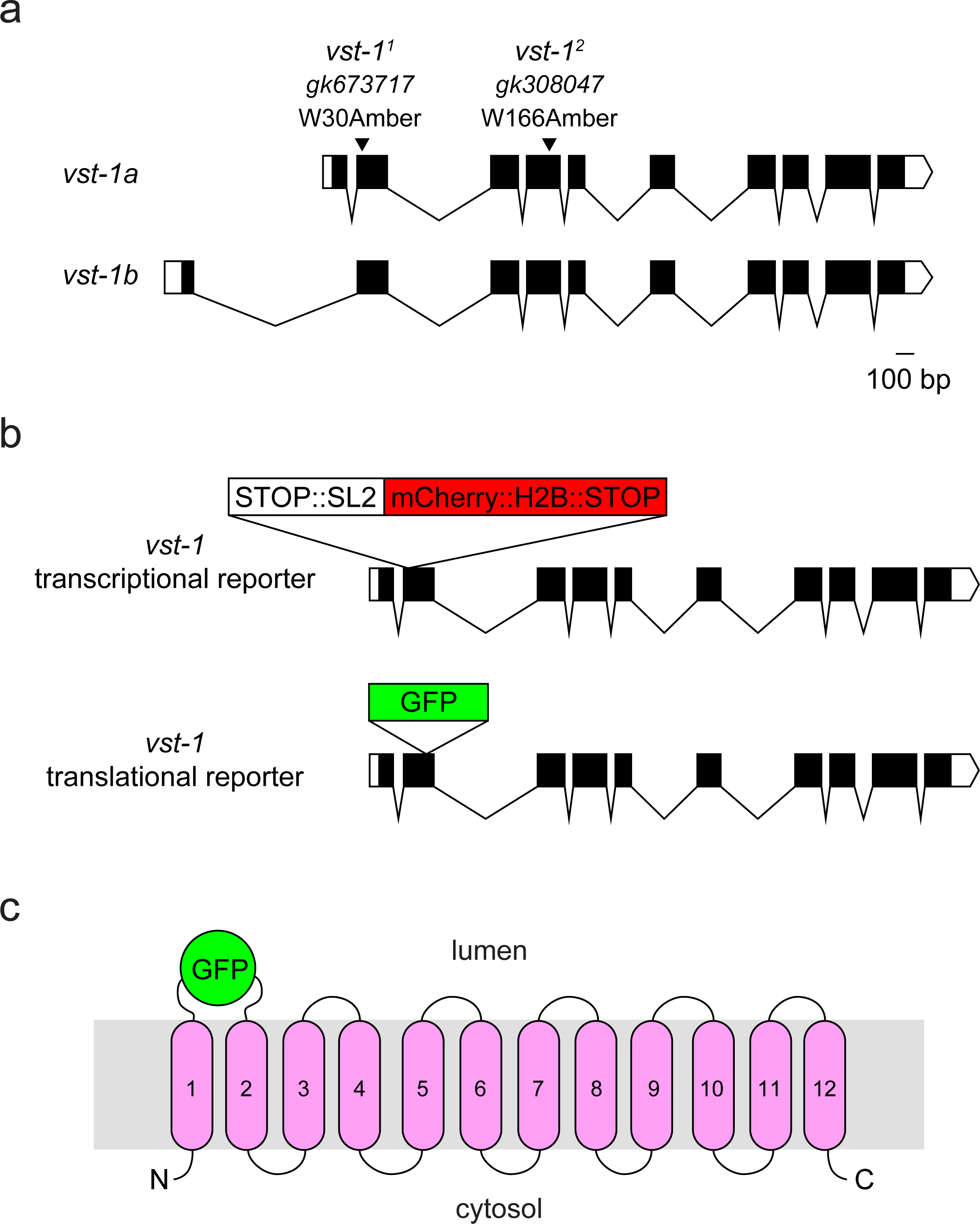
Alleles of *vst-1* and reporters of VST-1. (**a**) Alleles of *vst-1* used in this study. Arrowheads indicate locations of nonsense mutations of each allele in the second and fourth exons, respectively, of both isoforms of *vst-1*. (**b**) Schematic of the *vst-1* transcriptional fosmid reporter, *vst-1 fosmid::stop::SL2::1xNLS::mCherry::H2B::stop* (pEH62) and the *vst-1* translational reporter, *vst-1::gfp* (pJC106). (**c**) Schematic of the VST- 1::GFP fusion protein indicating the GFP insertion site within the first intralumenal loop.

**Supplemental Figure 2.**
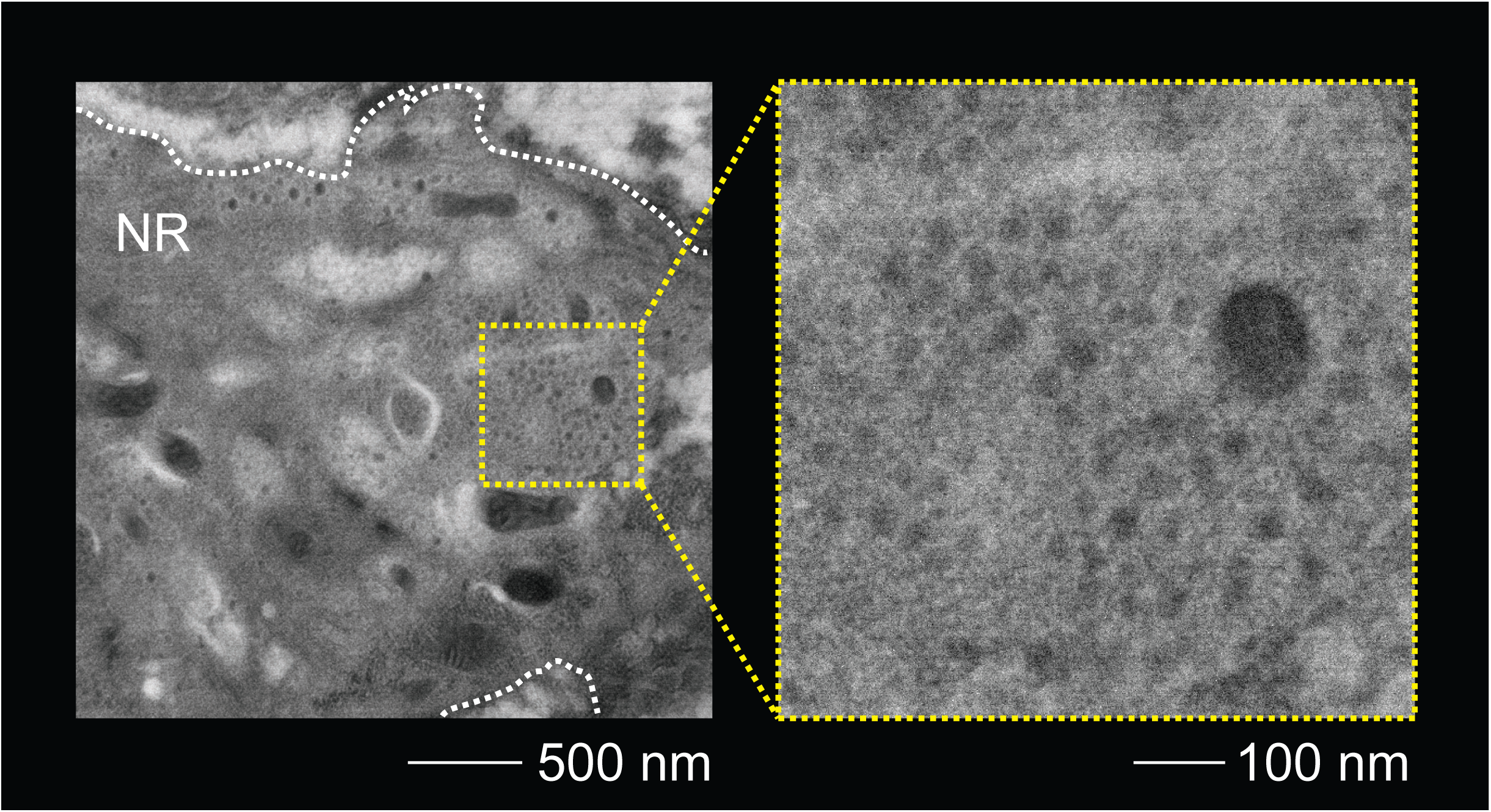
Electron micrograph of the nerve ring of a strain expressing VST- 1::GFP and stained with the PA15 secondary antibody only (no primary anti-GFP antibody). The left panel shows a larger field of view at a magnification of 31000x. The boundary of the nerve ring (NR) is marked by a dashed white line. The right panel shows a magnified view of the region indicated by dashed yellow lines. The dashed white line in the right panel indicates the perimeter of a single neurite cut in cross-section.

**Supplemental Figure 3.**
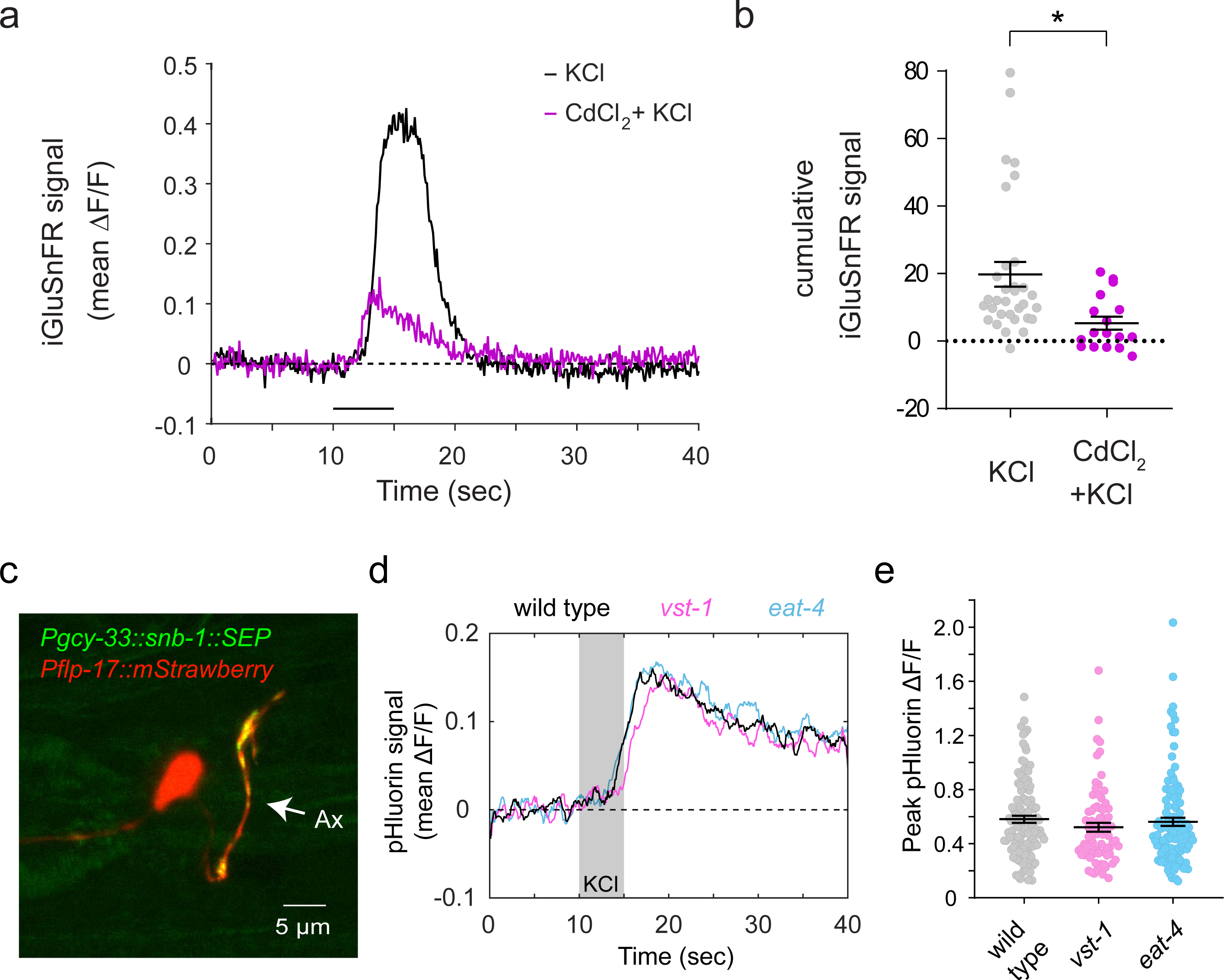
iGluSnFR signals recorded from wild-type BAG neurons in the presence of CdCl_2_ and synaptopHluorin signals recorded from wild-type, *eat-4*, and *vst-1* BAG neurons. (**a**) Mean iGluSnFR signal (mean ΔF/F) of cultured wild type BAG neurons stimulated by KCl in the presence and absence of CdCl_2_. The mean trace for KCl only was replotted from Figure 3a. (**b**) Cumulative iGluSnFR signal (area-under-the-curve of ΔF/F between 10-20 sec, mean ± SEM is shown) in the presence (*n* = 17) and absence (*n* = 32) of CdCl_2_. Asterisk indicates a statistical difference (p = 0.0012, Mann-Whitney test). (**c**) Micrograph showing the synaptopHluorin strain (*Pgcy-33::snb-1::superecliptic pHluorin; Pflp-17::mStrawberry; Punc-122::mCherry*) dissociated to obtain cultured BAG neurons. SynaptopHluorin expressed specifically in BAG neurons is enriched in puncta along the BAG axon indicating synaptic localization. (**d**) Plot shows mean synaptopHluorin signal (mean ΔF/F) of KCl-responsive puncta in response to 100 mM KCl after passing individual traces through a 1 sec moving-average filter. (**e**) Maximum synaptopHluorin signal (maximum ΔF/F between 10-20 sec) from KCl-responsive puncta from cultured wild type, *eat-4*, *vst-1* BAG neurons (n = 118, 115, 76, respectively). Quantification was conducted on the raw traces. There were no statistically significant differences among the genotypes (Kruskal-Wallis test, Dunn’s test for multiple comparisons between wild type and other genotypes).

**Supplemental Figure 4.**
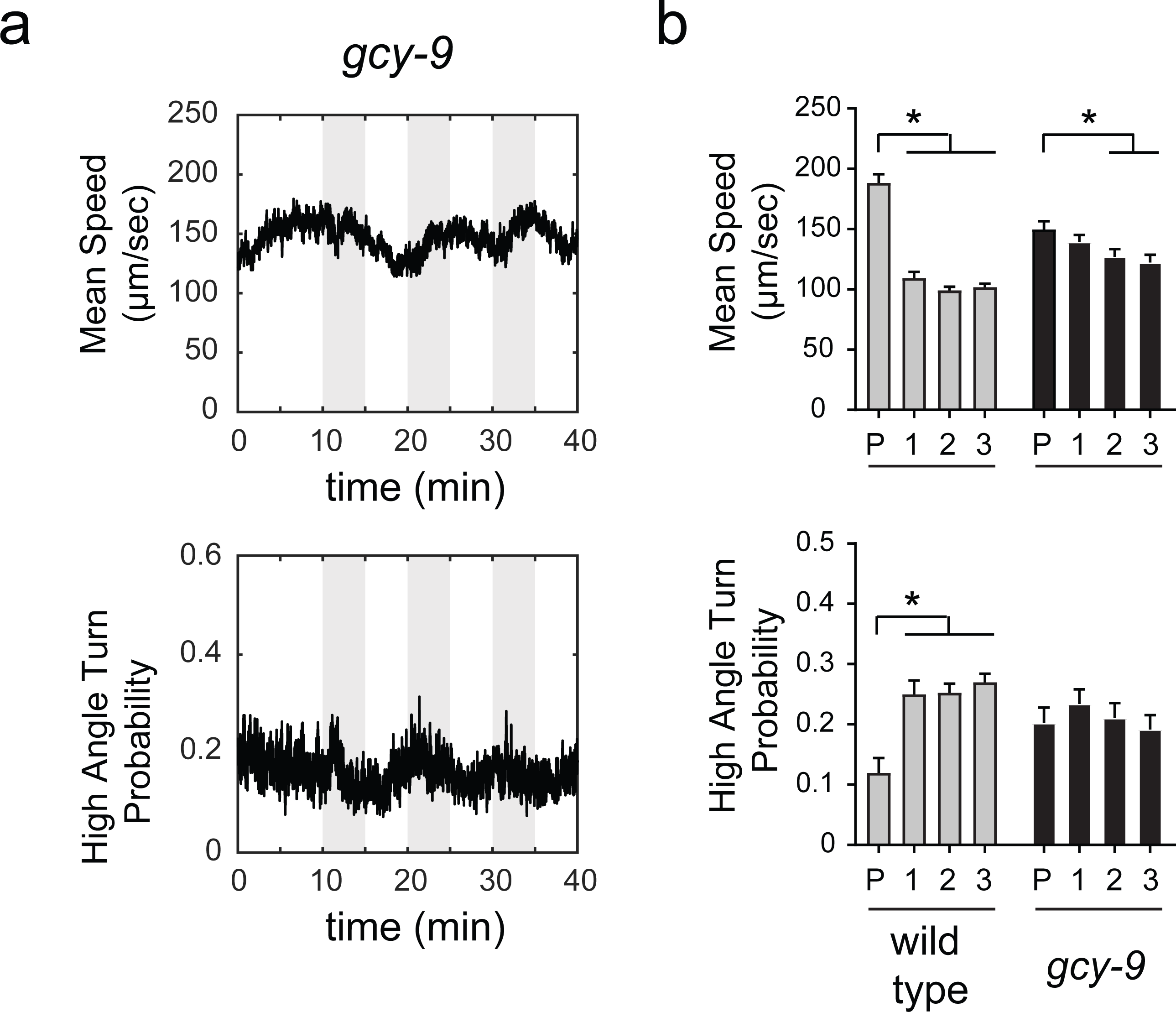
Analysis of locomotory behavior of *gcy-9* mutants. (**a**) Mean speed and high-angle turn probability changes over time for *gcy-9* mutants. Gray bars indicate CO_2_ pulses. (**b**) Mean speeds and turn probabilities ± SEM of wild type (replotted from Figure 5) and *gcy-9* mutants during the pre-stimulus period (P) and after each presentation of CO_2_ (labeled 1-3). The number of tracks analyzed (*n*) for mean speed during each period (P, 1, 2, and 3) are: 74, 105, 144, and 159 (wild type); 72, 70, 83, and 79 (*gcy-9*). The number of tracks analyzed (*n*) for high angle turn probability during each period (P, 1, 2, and 3) are: 74, 98, 145, and 158 (wild type); 72, 75, 80, and 80 (*gcy-9*). Asterisks indicate *p* < 0.05 for comparisons during each period to P using a Kruskal-Wallis test corrected for multiple comparisons via Dunn’s test (p-values for each comparison of mean speed marked with an asterisk – wild type P vs wild type 1: <0.0001; wild type P vs wild type 2: <0.0001; wild type P vs wild type 3: <0.0001; *gcy-9* P vs *gcy-9* 2: 0.022; *gcy-9* P vs *gcy-9* 3: 0.010; p-values for each comparison of high-angle turn probability marked with an asterisk - wild type P vs wild type 1: <0.0001; wild type P vs wild type 2: <0.0001; wild type P vs wild type 3: <0.0001).

**Supplemental Figure 5.**
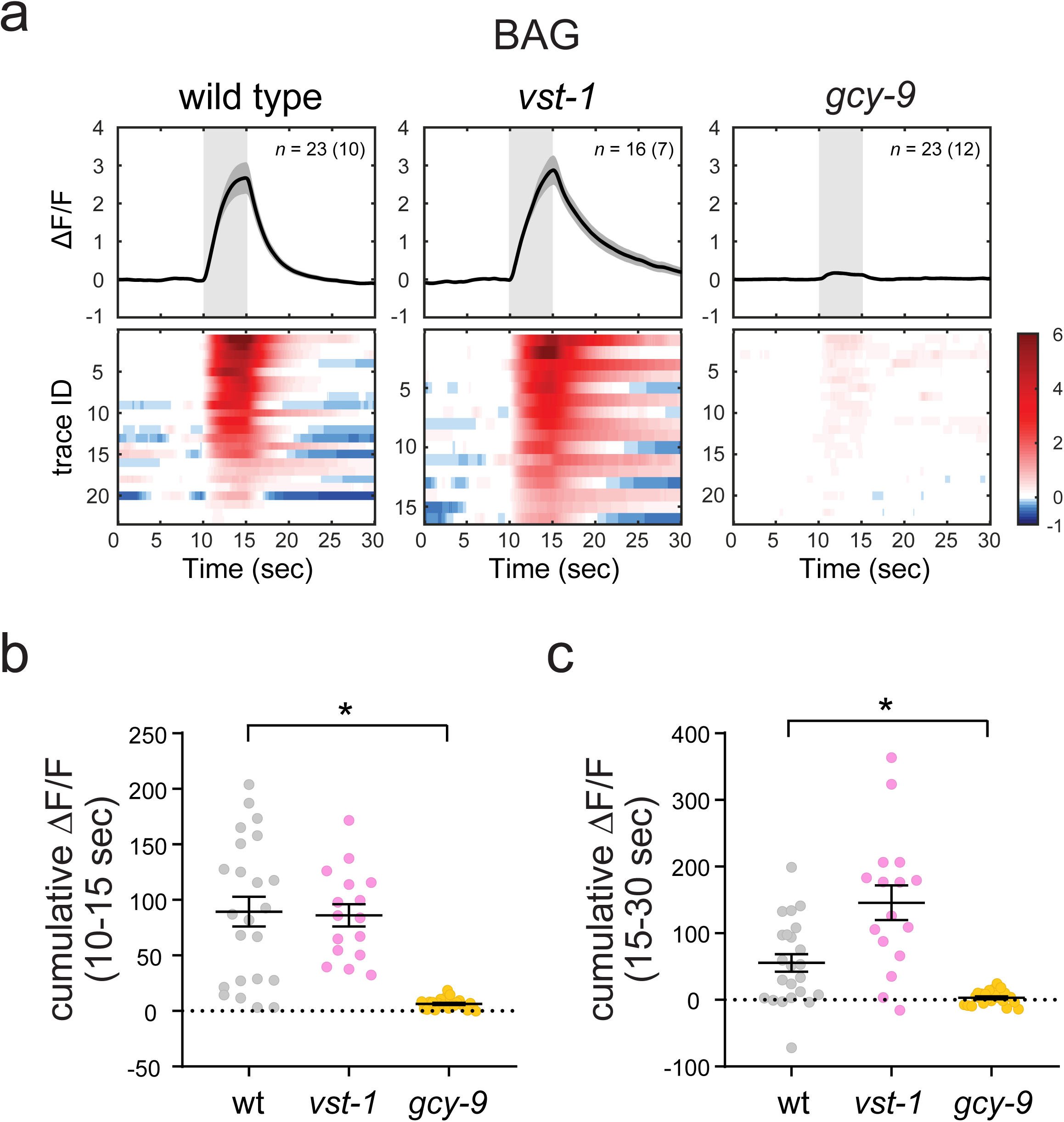
Calcium imaging of BAG neurons via microfluidic device. (**a**) Fluorescence changes of GCaMP6 expressed in BAG sensory neurons (*Pflp-17::GCaMP6f*) over time in wild-type, *vst-1*, and *gcy-9* animals. The number in parentheses indicates the number of worms analyzed and *n* indicates the total number of cells analyzed. Individual traces were passed through a 0.5 sec moving-average filter for the plots (raw traces were used for quantification of GCaMP signal in **b** and **c**). (**b**) Cumulative GCaMP signal between 10 and 15 seconds. Asterisk indicates p < 0.05 for comparisons of the mutants to wild type using a Kruskal-Wallis test corrected for multiple comparisons via Dunn’s test (p-value for comparison marked with an asterisk – wild type vs *gcy-9*: <0.0001). (**c**) Cumulative GCaMP signal between 15 and 30 seconds. Asterisks indicate p < 0.05 for comparisons of the mutants to wild type using a Kruskal-Wallis test corrected for multiple comparisons via Dunn’s test (p-value for comparison marked with an asterisk – wild type vs *gcy-9*: 0.0062). The p-value for wild type vs *vst-1* is 0.051.

**Supplemental Figure 6.**
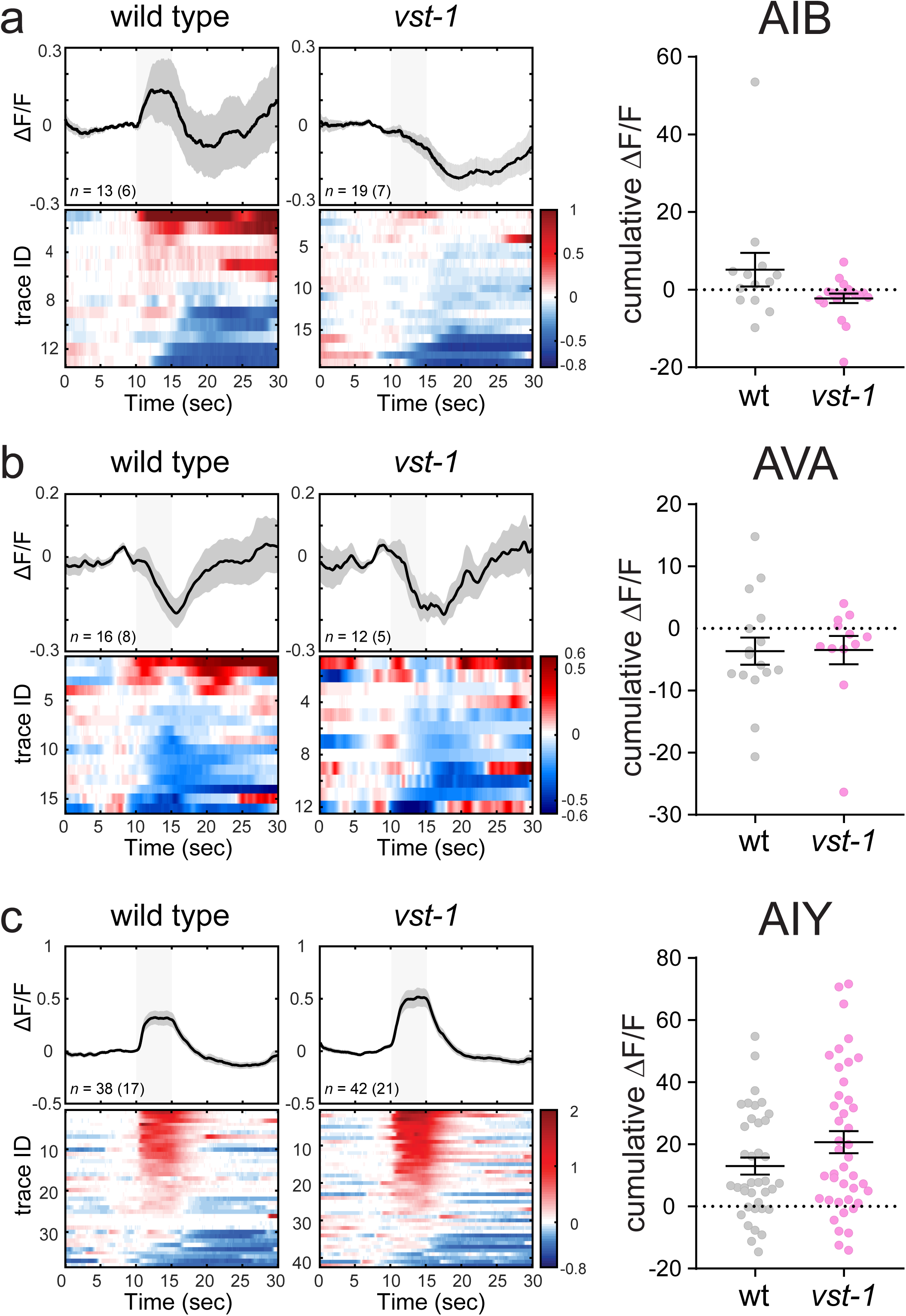
Calcium imaging of AIB, AVA, and AIY interneurons in the wild type and *vst-1* mutant backgrounds. (**a**) Plots of fluorescence changes of GCaMP3 in the soma of AIB interneurons of *Pinx- 1::GCaMP3* animals (left) and cumulative GCaMP signal (10-15 sec) (right). Statistical comparison between wild type and *vst-1* was conducted using a Mann-Whitney test (p = 0.071). (**b**)Plots of fluorescence changes of GCaMP6 in the soma of AVA interneurons of *Popt-3::GCaMP6f* animals (left) and cumulative GCaMP signal (10-15 sec) (right). Statistical comparison between wild type and *vst-1* was conducted using a Mann-Whitney test (p = 0.42). (**c**) Plots of fluorescence changes of GCaMP6 in the neurites of AIY interneurons of *Pttx- 3::GCaMP6s* animals (left) and cumulative GCaMP signal (10-15 sec) (right). Statistical comparison between wild type and *vst-1* was conducted using a Mann-Whitney test (p = 0.16). For **a-c**, individual traces were passed through a 0.5 sec moving-average filter for the plots (left) but the cumulative signal was quantified (right) using raw traces. The number in parentheses indicates the number of worms analyzed and *n* indicates the total number of cells analyzed.

**Supplemental Figure 7.**
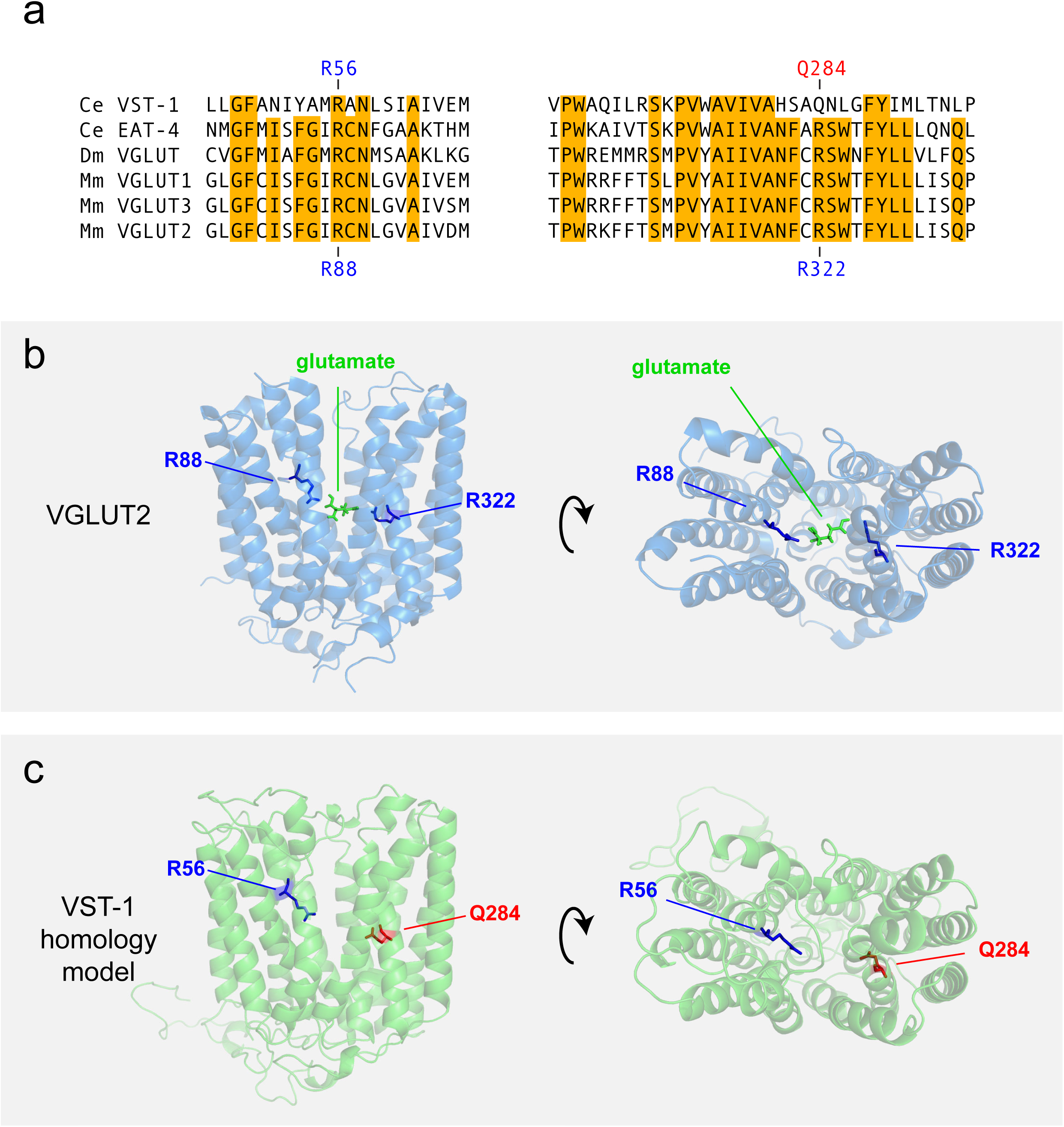
The transporter VST-1 lacks an arginine residue that is critical for glutamate-binding. (**a**) Sequence alignments of regions from the second and seventh transmembrane domains of VST-1 and VGLUTs of *C. elegans* (Ce), *D. melanogaster* (De), and *M. musculus* (Mm). Conserved residues are indicated in orange. Arginine 88 and Arginine 322 of VGLUT2, indicated in blue, are critical for VGLUT function and are thought to mediate glutamate-binding^49, 66, 67^. The corresponding residues in VST-1 are arginine 56 (blue) and glutamine 284 (red). (**b**) Structure of VGLUT2 and proposed interactions with glutamate after Li *et al.*^49^. Arginine 88 and arginine 322 are indicated in blue, and glutamate in green. (**c**) Homology model of VST-1 generated by Phyre2^68^ indicating conservation of one arginine required for glutamate transport (R56 in blue) and a substitution of glutamine for the other arginine (Q284 in red).

**Supplemental Figure 8.**
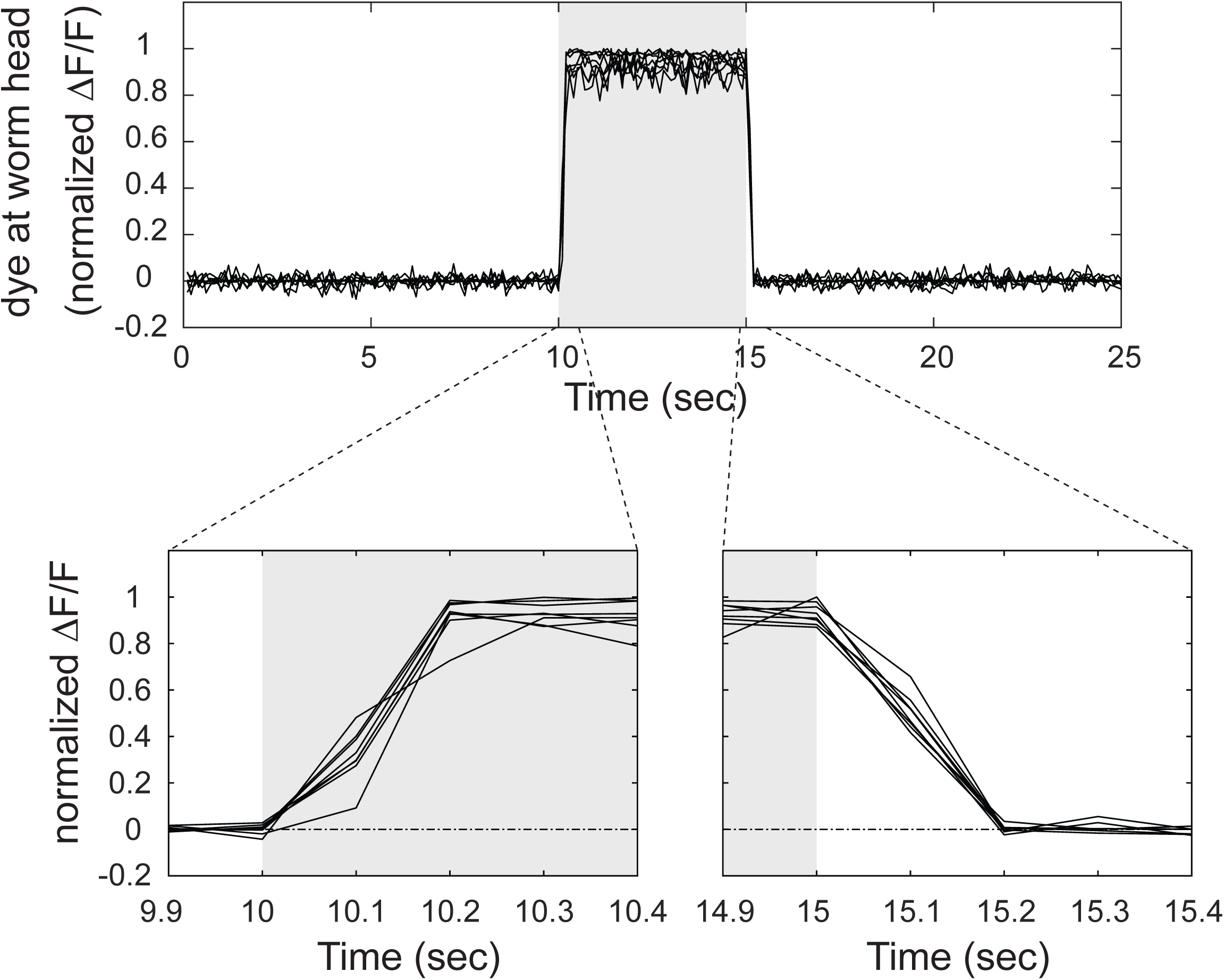
Microfluidic device driven by positive pressure can quickly and reliably switch between solutions. The upper panel shows change of fluorescein signal (ΔF/F) indicating the presence of solution in the channel directed toward the head of the worm (n = 8). The lower panels show the plot in the upper panel close to the time of valve switch. Gray indicates the window in which the head of the worm is exposed to fluorescein.

## SUPPLEMENTARY TABLES

**Supplementary Table 1.** **Statistical comparisons between different genotypes in** Figure 5. Statistical comparisons between all genotypes at corresponding periods (P,1,2,3) were conducted using a Kruskal-Wallis test corrected for multiple comparisons by Dunn’s test and the p-values are indicated for each comparison. Bold numbers indicate p < 0.05.

Mean Speed
| Comparisons | Period |  |  |  |
| --- | --- | --- | --- | --- |
|  | Pre-stimulus | Stimulus 1 | Stimulus 2 | Stimulus 3 |
| wild type vs *vst-1* | **<0.0001** | **<0.0001** | **<0.0001** | **<0.0001** |
| wild type vs *eat-4* | **<0.0001** | **<0.0001** | **<0.0001** | **<0.0001** |
| wild type vs *eat-4; vst-1* | **<0.0001** | **0.0141** | **<0.0001** | **0.0020** |
| wild type vs *glr-1* | >0.9999 | >0.9999 | >0.9999 | >0.9999 |
| wild type vs *glr-1; vst-1* | >0.9999 | 0.1834 | >0.9999 | >0.9999 |
| *vst-1* vs *eat-4* | >0.9999 | **0.0134** | **<0.0001** | **<0.0001** |
| *vst-1* vs *eat-4; vst-1* | >0.9999 | **0.0004** | **<0.0001** | **<0.0001** |
| *vst-1* vs *glr-1* | **<0.0001** | **<0.0001** | **<0.0001** | **<0.0001** |
| *vst-1* vs *glr-1; vst-1* | **<0.0001** | **<0.0001** | **<0.0001** | **<0.0001** |
| *eat-4* vs *eat-4; vst-1* | >0.9999 | >0.9999 | >0.9999 | 0.2952 |
| *eat-4* vs *glr-1* | **<0.0001** | **0.0138** | **<0.0001** | **<0.0001** |
| *eat-4* vs *glr-1; vst-1* | **<0.0001** | **<0.0001** | **<0.0001** | **<0.0001** |
| *eat-4; vst-1* vs *glr-1* | **<0.0001** | 0.4878 | **<0.0001** | **0.0204** |
| *eat-4; vst-1* vs *glr-1; vst-1* | **<0.0001** | **<0.0001** | **<0.0001** | **0.0014** |
| *glr-1* vs *glr-1; vst-1* | >0.9999 | **0.0164** | >0.9999 | >0.9999 |

Turn Probability
| Comparisons | Period |  |  |  |
| --- | --- | --- | --- | --- |
|  | Pre-stimulus | Stimulus 1 | Stimulus 2 | Stimulus 3 |
| wild type vs *vst-1* | >0.9999 | >0.9999 | **0.0013** | **0.0011** |
| wild type vs *eat-4* | >0.9999 | >0.9999 | >0.9999 | **0.0167** |
| wild type vs *eat-4*; *vst-1* | >0.9999 | >0.9999 | **0.0002** | **<0.0001** |
| wild type vs *glr-1* | 0.6504 | >0.9999 | 0.0620 | **0.0092** |
| wild type vs *glr-1*; *vst-1* | **0.0416** | 0.3433 | **0.0007** | 0.1508 |
| *vst-1* vs *eat-4* | >0.9999 | >0.9999 | **<0.0001** | **<0.0001** |
| *vst-1* vs *eat-4*; *vst-1* | >0.9999 | 0.7648 | **<0.0001** | **<0.0001** |
| *vst-1* vs *glr-1* | >0.9999 | >0.9999 | **<0.0001** | **<0.0001** |
| *vst-1* vs *glr-1*; *vst-1* | 0.4635 | 0.1718 | **<0.0001** | **<0.0001** |
| *eat-4* vs *eat-4*; *vst-1* | >0.9999 | **0.0384** | 0.0670 | **0.0355** |
| *eat-4* vs *glr-1* | **0.0458** | 0.9114 | >0.9999 | >0.9999 |
| *eat-4* vs *glr-1*; *vst-1* | **0.0006** | **0.0063** | 0.1482 | >0.9999 |
| *eat-4*; *vst-1* vs *glr-1* | >0.9999 | >0.9999 | >0.9999 | 0.5437 |
| *eat-4*; *vst-1* vs *glr-1*; *vst-1* | 0.2334 | >0.9999 | >0.9999 | **0.0164** |
| *glr-1* vs *glr-1*; *vst-1* | >0.9999 | >0.9999 | >0.9999 | >0.9999 |

**Supplementary Table 2.** **Mutant and transgenic lines used for this study.**

| Strain | Genotype |
| --- | --- |
| FQ306 | *gcy-9(tm2816)* |
| MT6308 | *eat-4(ky5)* |
| VC40514 | *vst-1(gk673717)* |
| VC20397 | *vst-1(gk308047)* |
| FQ1151 | *wzEx308[Pflp-17::eat-4 RNAi; Punc-122::mCherry]* |
| FQ1160 | *wzEx315[Pflp-17::vst-1 RNAi; Punc-122::GFP]* |
| FQ1571 | *otIs388[eat-4 fosmid::SL2::YFP::H2B + (pBX) pha-1(+)] pha-1(e2123); wzEx431[vst-1 fosmid::stop::SL2::1xNLS::mCherry::H2B; Punc-122::GFP]* |
| FQ1137 | *vst-1(gk308047); wzEx305[vst-1::GFP; Pflp-17::mStrawberry]* |
| FQ2656 | *unc-104; vst-1(gk308047); wzEx305 [vst-1::GFP; Pflp-17::mStrawberry]* |
| FQ1167 | *wzEx323[vst-1::GFP; eat-4::mCherry; Punc-122::GFP]* |
| FQ843 | *wzEx204[Pflp-17::iGluSnFR; Punc-122::mCherry]* |
| FQ891 | *eat-4(ky5); wzEx204[Pflp-17::iGluSnFR; Punc-122::mCherry]* |
| FQ1174 | *vst-1(gk673717); wzEx204[Pflp-17::iGluSnFR; Punc-122::mCherry]* |
| FQ911 | *vst-1(gk308047); wzEx204[Pflp-17::iGluSnFR; Punc-122::mCherry]* |
| FQ1575 | *wzEx434[Pgcy-33::snb-1::superecliptic pHluorin; Pflp-17::mStrawberry; Punc-122::mCherry]* |
| FQ1764 | *vst-1(gk308047); wzEx434[Pgcy-33::snb-1::superecliptic pHluorin; Pflp-17::mStrawberry; Punc-122::mCherry]* |
| FQ1817 | *eat-4(ky5); wzEx434[Pgcy-33::snb-1::superecliptic pHluorin; Pflp-17::mStrawberry; Punc-122::mCherry]* |
| KP4 | *glr-1(n2461)* |
| FQ1277 | *glr-1(n2461); vst-1(gk308047)* |
| FQ1440 | *eat-4(ky5); vst-1(gk308047)* |
| FQ845 | *wzEx165[Pflp-17::GCaMP6f; Punc-122::mCherry]* |
| FQ2110 | *vst-1(gk308047); wzEx165[Pflp-17::GCaMP6f; Punc-122::mCherry]* |
| FQ2143 | *gcy-9(tm2816); wzEx165[Pflp-17::GCaMP6f; Punc-122::mCherry]* |
| CX13440 | *kyEx4018[Pinx-1::GCaMP3; Punc-122::dsRed]* |
| FQ2039 | *vst-1(gk308047); kyEx4018[Pinx-1::GCaMP3; Punc-122::dsRed]* |
| FQ944 | *wzEx246[Popt-3::GCaMP6f; Punc-122::mCherry]* |
| FQ1922 | *vst-1(gk308047); wzEx246[Popt-3::GCaMP6f; Punc-122::mCherry]* |
| FQ2348 | *sraEx490[Pttx-3::GCaMP6s]* |
| FQ2236 | *vst-1(gk308047); sraEx490[Pttx-3::GCaMP6s]* |
| ZC1508 | *yxIs19[Pglr-3a::GCaMP3; Punc-122::dsRed]* |
| FQ2040 | *yxIs19[Pglr-3a::GCaMP3; Punc-122::dsRed]; vst-1(gk308047)* |
| FQ492 | *yxIs19[Pglr-3a::GCaMP3; Punc-122::dsRed]; gcy-9(tm2816)* |
| FQ2178 | *yxIs19[Pglr-3a::GCaMP3; Punc-122::dsRed]; gcy-9(tm2816) vst-1(gk308047)* |
| FQ2176 | *glr-1(n2461); yxIs19[Pglr-3a::GCaMP3; Punc-122::dsRed]* |
| FQ2177 | *glr-1(n2461); yxIs19[Pglr-3a::GCaMP3; Punc-122::dsRed]; vst-1(gk308047)* |
| TV2217 | *wyIs93[Pglr-3::mCherry::rab-3; Pglr-3::glr-1::GFP; Punc-122::RFP]; wyEx828[Pglr-3::caspase-3(p12)::nz; Pglr-3::cz::caspase-3(p17); Pglr-3::mCherry; Punc-122::GFP]* |
| FQ2476 | *vst-1(gk308047) wyIs93[Pglr-3::mCherry::rab-3; Pglr-3::glr-1::GFP; Punc-122::RFP]; wyEx828[Pglr-3::caspase-3(p12)::nz; Pglr-3::cz::caspase-3(p17); Pglr-3::mCherry; Punc-122::GFP]* |

**Supplementary Table 3.**
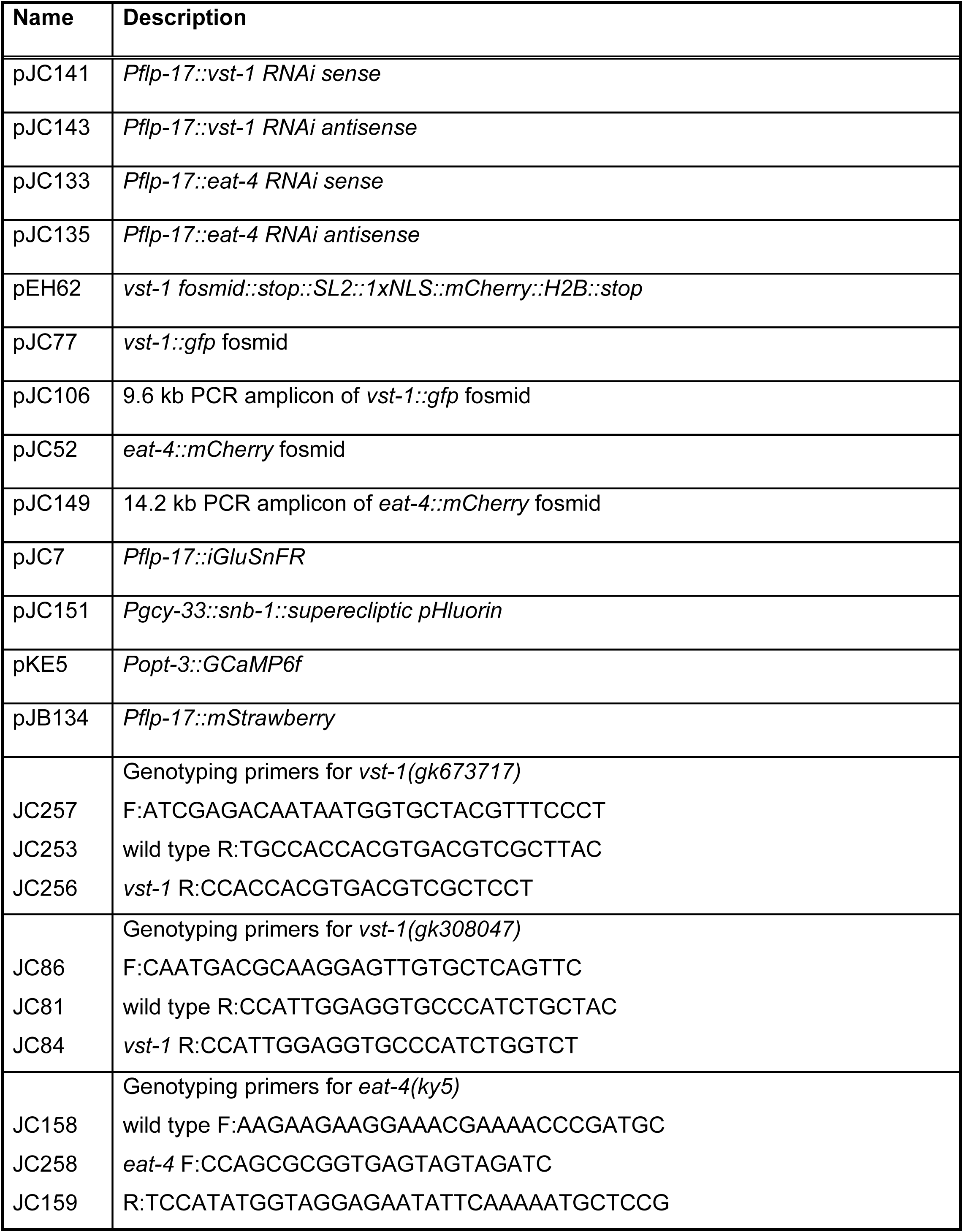
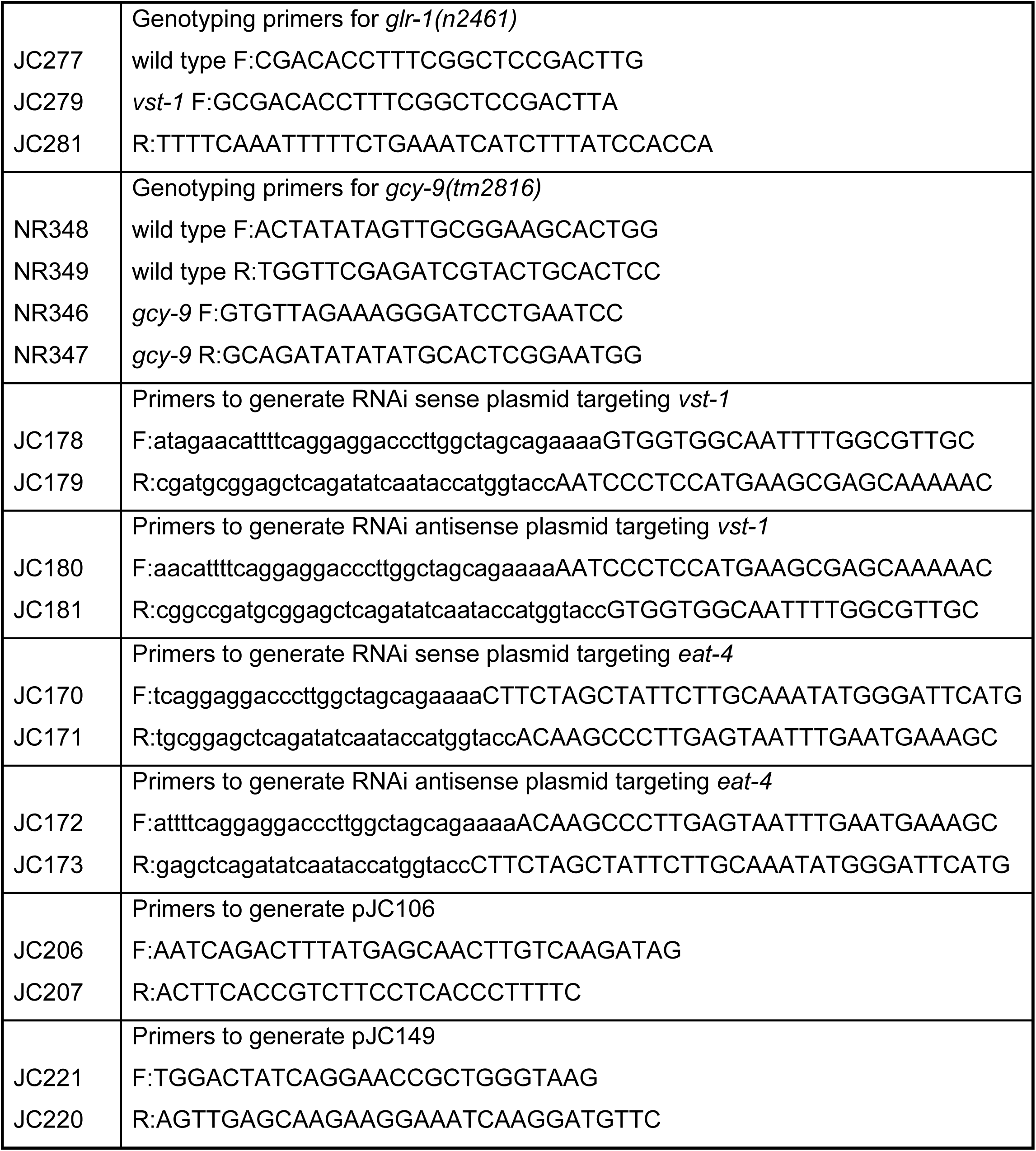
**Plasmids, fosmids, and primers used for this study.**

